# Balanced mitochondrial and cytosolic translatomes underlie the biogenesis of human respiratory complexes

**DOI:** 10.1101/2021.05.31.446345

**Authors:** Iliana Soto, Mary Couvillion, Erik McShane, Katja G. Hansen, J. Conor Moran, Antoni Barrientos, L. Stirling Churchman

## Abstract

Oxidative phosphorylation (OXPHOS) complexes consist of nuclear and mitochondrial DNA-encoded subunits. Their biogenesis requires cross-compartment gene regulation to mitigate the accumulation of disproportionate subunits. To determine how human cells coordinate mitochondrial and nuclear gene expression processes, we established an optimized ribosome profiling approach tailored for the unique features of the human mitoribosome. Analysis of ribosome footprints in five cell types revealed that average mitochondrial synthesis rates corresponded precisely to cytosolic rates across OXPHOS complexes. Balanced mitochondrial and cytosolic synthesis did not rely on rapid feedback between the two translation systems. Rather, *LRPPRC*, a gene associated with Leigh’s syndrome, is required for the reciprocal translatomes and maintains cellular proteostasis. Based on our findings, we propose that human mitonuclear balance is enabled by matching OXPHOS subunit synthesis rates across cellular compartments, which may represent a vulnerability for cellular proteostasis.

---

Of the >1000 proteins that localize to human mitochondria (*1*), only 13 are encoded on mitochondrial DNA (mtDNA) and synthesized by mitochondrial ribosomes (mitoribosomes) in the mitochondrial matrix. These proteins, which comprise the core subunits of the oxidative phosphorylation (OXPHOS) complexes, are co-assembled with >45 nuclear DNA (nDNA) encoded OXPHOS subunits that are synthesized by cytosolic ribosomes (cytoribosomes) and imported into mitochondria. The coordinated expression of the nuclear and mitochondrial genomes maintains mitonuclear balance, in which mtDNA- and nDNA-encoded OXPHOS subunits are in equilibrium, ensuring accurate assembly and proper function of complexes (*2–4*). Unassembled subunits can overwhelm proteostasis pathways and assembly intermediates are at risk of producing reactive oxygen species (ROS); both of these consequences are detrimental to the cell and lead to disease and aging phenotypes (*5–9*). In *S. cerevisiae*, cytosolic and mitochondrial translation programs occur synchronously during OXPHOS biogenesis (*10*). Thus, cross-compartment translation regulation (*9*) contributes to mitonuclear balance in yeast cells.

Expression and regulation of mtDNA differ substantially between fungal and animal cells. Yeast mt-mRNAs have long 5’ leaders (up to ∼1 kb) to which nuclear DNA-encoded translation activators (TA) bind to recruit mitoribosomes to particular mt-mRNAs, enabling nuclear gene expression to directly control mitochondrial translation (*11–13*). However, analogous mechanisms of translational control are not thought to exist in human cells. Human mtDNA molecules are transcribed as two polycistronic transcripts that are processed largely through the excision of mt-tRNAs flanking protein-coding mt-mRNAs, leaving most mt-mRNAs leaderless. In addition, most yeast TAs are not conserved in humans, and those that are serve different functions (*14*). To date, only one TA for human mitochondrial translation has been identified (*15*). Furthermore, in human cells, protein quality control pathways rapidly degrade superstoichiometric nuclear DNA-encoded OXPHOS subunits (*4, 9, 16*), and multiple transcriptional programs adapt mitochondrial activity to environmental perturbations (*17, 18*). Thus, human and yeast cells may use different strategies to balance mitonuclear expression and OXPHOS assembly.

In this study, we quantify mitoribosome density across all transcripts with subcodon resolution in five human cell types. The resultant detailed view of translation across the 13 canonical open reading frames (ORF) revealed features of translation initiation on leaderless mt-mRNAs and a downstream ORF in the 3’ UTR of *MT-ND5*. Remarkably, we find that average synthesis of nDNA- and mtDNA-encoded OXPHOS subunits for each complex were highly correlated in all cell types analyzed. In contrast to the situation in yeast, correlated translatomes in human cells were not achieved through rapid crosstalk between cytosolic and mitochondrial translation systems. Instead, we find that the nDNA-encoded protein LRPPRC, which stabilizes most mt-RNAs and promotes their polyadenylation and loading onto the mitoribosome (*19–23*), was required to balance mitochondrial and cytosolic protein synthesis.

## Results

### High-resolution profiling of human mitochondrial translation

The human mitoribosome is highly proteinaceous, containing twice as much protein as RNA, and is largely associated with the inner membrane in order to facilitate co-translational insertion of OXPHOS proteins (*24*). These unique features make quantitative biochemical purification of mitoribosomes for the purpose of analyzing their translatomes more difficult than purification of cytosolic ribosomes (*25*). To dissect mitochondrial translation with high resolution and minimum bias, we re-engineered ribosome profiling to accommodate the unique characteristics of the human mitoribosome, refining each step of the approach (**Fig 1A)**.

**Figure 1.**
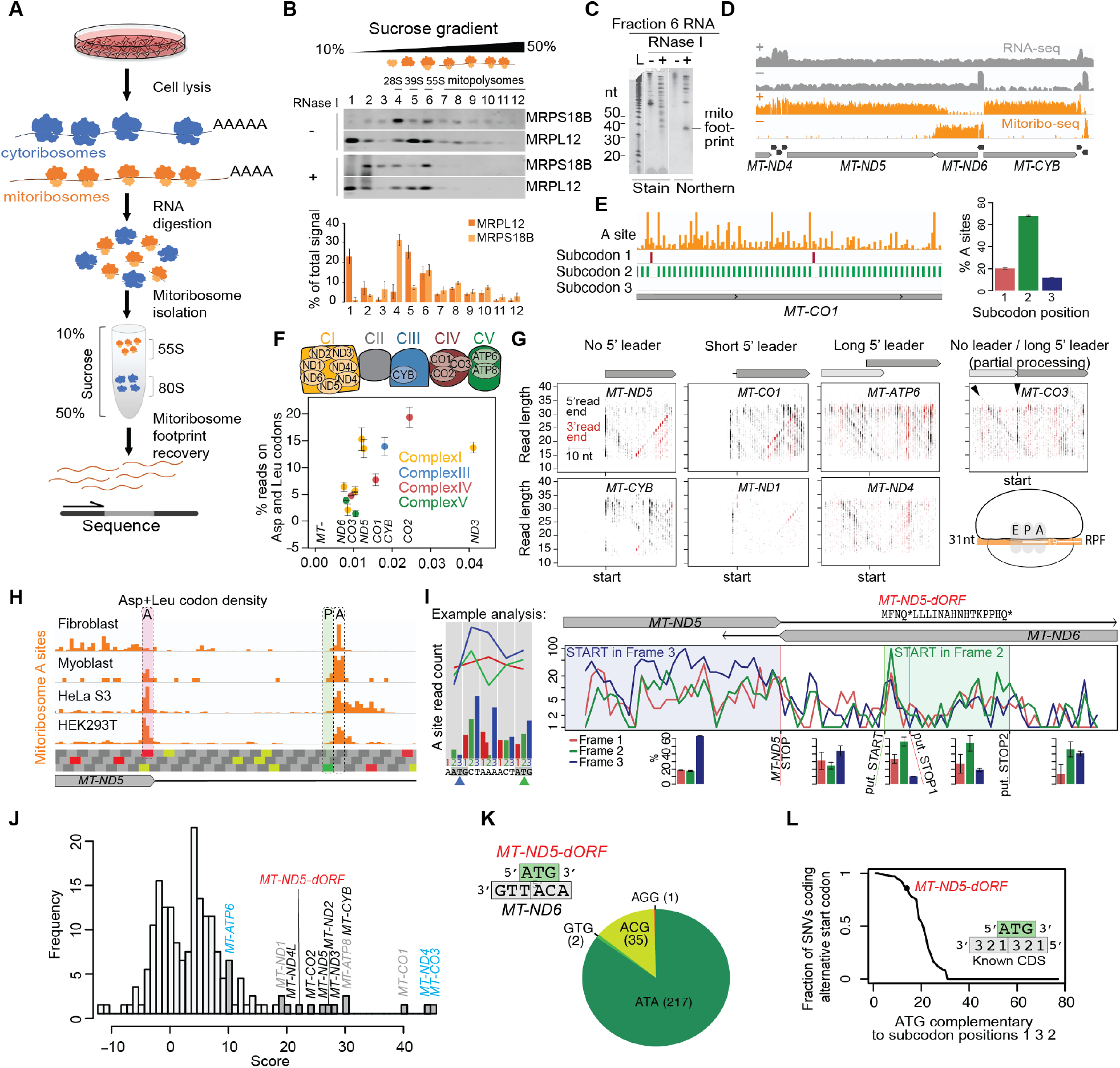
Human mitoribosome profiling and resolved features of mitochondrial translation. (**A**) Schematic of the human mitoribosome profiling approach. (**B**) Isolation of mitomonosome (55S) footprints from mitochondrial polysomes. Digested (+RNase I) and undigested (-RNase I) HeLa S3 lysates were clarified and loaded onto linear 10-50% sucrose gradients. Detection of mitoribosomes in each fraction (top panel) was achieved by western blots against proteins of the large (MRPL12) and small subunit (MRPS18B). Signals of both subunits from four experiments were quantified, showing a periodic trend (bottom panel). (**C**) Footprints recovered from the mitomonosome fraction (Fraction 6) were subjected to northern blot (*MT-CO2* probe) for assessment of size distribution. (**D**) Sequencing data over a representative portion of the mitochondrial genome, with mRNAs (labeled) and tRNAs (black) annotated below. Panels show RNA-seq read coverage (gray) and mitoribosome profiling (mitoribo-seq) read A-site density (orange) on heavy (+) and light (-) strands. Y axis is in log scale. (**E**) Left panel: A-site density (orange) across a portion of *MT-CO1* with subcodon position of maximum signal plotted below. Right panel: Percentage of reads with their A sites on each subcodon position across all mt-mRNAs. (**F**) Percentage of reads on each gene (color coded by their location in OXPHOS complexes) on Asp and Leu (UUR) codons. Genes are ordered along the x axis by their Asp and Leu (UUR) codon density with select genes labeled. Error bars show range across two fibroblast replicates. (**G**) Scatterplots to visualize mitoribosome position during initiation. Start-codon-aligned RPF 5’ and 3’ ends are plotted according to their genome position and read length. Arrowheads highlight *MT-CO3* initiation on both unprocessed *MT-ATP8/6-CO3* and processed *MT-CO3* mRNA. Cartoon shows positioning of mitoribosome E, P, and A sites on a full-length (31 nt) RPF. Short 5’ leaders consist of one to three nts. (**H**) Translation initiation occurs on an internal AUG in the *MT-ND5* 3’ UTR across cell lines. A-site density on the *MT-ND5* stop codon (shaded in red) and P-site density (shaded in green, A-site density one codon downstream) of initiating ribosomes on the internal AUG are highlighted. Codons in the three frames are shown below with stop (UAA, UAG), start (AUG), and mitochondrial alternative start (AUU, AUA) highlighted (red, green, yellow, respectively). (**I**) Stacked frame plot highlighting number of A-site transformed reads in each subcodon position on heavy strand. The example analysis (left panel) shows how raw A site read counts are transformed to line plots. Mitoribosome profiling data are from two fibroblast replicates, summed. Amino acid sequence of putative MT-ND5-dORF peptide is shown with putative (put.) start and stop codons highlighted with green and red lines, respectively. Bar plots below show the average percentage of reads in each subcodon position for regions indicated, with error bars showing range between the replicates summed above. (**J**) Distribution of initiation scores after applying ad hoc scoring method to fibroblasts. Gene names in black: known genes with no leader; gray: known genes with 1 to 3 nt leader; cyan: known genes with long 5’ UTR. (**K**) Variants at *MT-ND5-dORF* ATG (in green), which is antisense to *MT-ND6* (in grey) in the orientation indicated by the schematic. ATA: canonical alternative mitochondrial start codon. GTG: variant start codon shown to efficiently initiate translation. ACG: capable of initiation in reconstituted mammalian mitochondria translation (*37*). AGG: ambiguous (stop or frame-shift inducing). (**L**) Genome-wide analysis of SNVs at ATGs. Only equivalently oriented ATGs opposite to codons in known ORFs were included to equalize the constraints (see methods). ATGs were required to have at least 2 SNVs to be included.

We first identified lysis and purification conditions that improved the recovery of mitoribosomes (**Fig S1A**). The 55S mitoribosome sediments separately from the 80S cytoribosome on a linear sucrose gradient (**Fig S1B**). Because RNA absorbance measurements are dominated by cytoribosomes, we used western blot analysis of both the small and large mitoribosome subunits to follow mitoribosome sedimentation (**Fig 1B**). We observed that both mitoribosome subunits peaked in a single monosome fraction, followed by a periodic co-sedimentation through later fractions, indicative of polysomes, which have been challenging to capture previously (*26, 27*). The polysomes and monosomes collapsed in the presence of EDTA (**Fig S1B**), resulting in differential sedimentation of each subunit. We did not observe polysomes under non-optimized conditions (data not shown). Thus, our optimized lysis enriches for mitochondrial polysomes (**Fig 1B**), permitting a broader view of mitochondrial translation.

Careful titration of RNase treatment revealed a mitoribosome footprint of 31–34 nucleotides (nt) (**Fig 1C, Fig S1C**). We analyzed mitoribosome footprints in five human cell types: primary fibroblasts, primary myoblasts, myocytes (post-differentiation-induction day 2 myoblasts), HeLa S3 cells, and HEK293T cells (**Fig S1D**). Sequencing and alignment of mitoribosome footprints revealed specific coverage of mitochondrial coding sequences (**Fig 1D**). Our optimized approach led to ∼10x more reads aligning to mitochondrial mRNAs compared to other protocols (**Fig S1E, Table S1**). Importantly, we achieved higher percentages of codons covered and stronger three-nucleotide periodicity (**Fig 1E, Table S1**). We also purified and sequenced footprints from mitoribosomes that had been further purified after sedimentation by immunoprecipitation (**Fig S1F**), which increased the fraction of reads mapped to mitochondrial mRNA but did not impact data quality (**Table S1**). Finally, we performed mitoribosome profiling in the presence or absence of cytosolic and mitochondrial translation inhibitors and obtained comparable results under both conditions (**Fig. S1G**). All subsequent experiments were performed without translation inhibitors and without immunoprecipitation.

Due to the high resolution of our re-engineered approach, we were able to detect mitoribosome pausing on all codons across the cell lines, including canonical and non-canonical termination codons (**Fig S2A**). Pausing at aspartate (Asp) and leucine (Leu) was pervasive, with mitoribosome stalling observed at nearly every occurrence of either codon (**Fig S2B**). In myoblasts, where tRNA abundance data are available, codon occupancy did not correlate with tRNA abundance, and the abundance of tRNA^Asp^ was above average (**Fig S2C**) (*28*). The malate– aspartate shuttle transfers Asp out of the mitochondrial matrix to the cytosol (**Fig S2D**); consequently, the matrix concentration of Asp is ∼7x lower than that of glutamate (*29*). This may explain the high mitoribosome occupancy on Asp codons. The high mitoribosome density at Asp and Leu codons is substantial; ribosome density on these codons constitute up to 20% of the total density for some genes (**Fig 1F**). These observations hint at a link between mitochondrial translation and metabolic pathways.

### Resolved features of mitochondrial translation

To analyze mitochondrial translation initiation on mRNAs with minimal or no 5’ UTRs, we visualized mitoribosome positions during initiation by analyzing ribosome-protected fragment (RPF) position versus RPF size (**Fig 1G**). *MT-ATP6* and *MT-ND4* mRNAs have long leaders, so the high ribosome density at the start codon is visualized as a V, due to some reads arising from over- or under-digested RPFs. For mRNAs with a concise (1–3 nt) leader or no (0-nt) leader, the plots reveal an enrichment of 15-nt half-size RPFs increasing in size to full-length, at which point the mRNA channel is fully occupied. These shorter half-footprints have not been previously detected (*30*) and indicate that mitoribosomes load at the ends of messages with the P site on the start codon. Translation initiation occurs from the canonical AUG but also AUA and AUU. Initiation from non-canonical mitochondrial start codons has been proposed to be less efficient (*31*); however, we did not observe this trend in the cell types that we analyzed (**Fig S2E**).

In general, the length of the 5’ leader did not correlate with translation efficiency (**Fig S2E**), with one exception. Most mitochondrial mRNAs are rapidly excised from the long nascent polycistronic transcript (*32*). However, unprocessed *ATP8/ATP6-CO3* mRNA accumulates to higher levels than other processing intermediates (*33*) and we observed *MT-CO3* translation initiation on both processed and unprocessed transcripts (**Fig 1G, Fig S2F**). The frequency of *MT-CO3* mRNA translation initiation varied across cell types but was up to 4.3-fold higher on unprocessed transcripts (**Fig S2F**). Thus, for *MT*-*CO3* mRNA, the presence of a leader sequence favors translation initiation.

### Mitoribosome-engaged noncanonical open reading frame

Across all cell lines analyzed, we observed a peak of ribosome density in the *MT-ND5* 3’ UTR. The density starts with high A-site occupancy one codon downstream of an AUG initiation codon (**Fig 1H**), characteristic of an initiating ribosome with its P site on the start codon. The AUG begins a short ORF encoding a four–amino acid putative peptide (**Fig 1I, S3A**) that ends with a stop codon followed by a C, a context that promotes read-through in the cytosol (*34*). Because in-frame RPFs were detected beyond the stop codon (**Fig 1I, S3A**), the density could indicate the production of a four amino acid or longer peptide encoded in the *MT-ND5* 3’ UTR. Other mitochondrial small ORFs (smORFs) encode peptides, most prominently humanin and MOTS-c, encoded within the mitochondrial rRNA genes (*35*). It is not known whether these smORFs are translated in the mitochondria or cytosol. We found no evidence for their translation by mitoribosomes (**Fig S3B**), although rRNA is a prominent contaminant in ribosome profiling data and could obscure footprint periodicity patterns.

To determine whether the *MT-ND5* downstream ORF (*MT-ND5-dORF*) is likely to be translated, we devised a strategy to score all start codons across the genome using mitoribosome occupancy. Three features were integrated into the scoring design: (1) P-site pausing, (2) periodicity/in frame signal, and (3) overall ribosome occupancy before and after the start codon (see Methods for details; **Fig S3C**). Start codons of known genes all scored highly, regardless of whether the ORF they began had no leader, a short leader, or a long 5’ UTR as in overlapping ORFs (**Fig 1J**). *MT-ND5-dORF* scored amongst the known genes across cell lines (**Fig 1J, S3D**), suggesting that it is translated. Mitoribosomes density across *MT-ND5-dORF* indicate that it is translated at levels comparable to mtDNA-encoded OXPHOS Complex I subunits (**Fig. S3E**).

We next investigated whether *MT-ND5-dORF* resists accumulating mutations. We analyzed a catalog of mtDNA variants across a population of nearly 200,000 individuals not enriched for mitochondrial disorders (*36*). Of 255 single-nucleotide variants (SNVs) of the *MT-ND5-dORF* AUG start codon, 219 produce a known alternative mitochondrial start codon (**Fig 1K**). Of the 36 that do not, 35 result in ACG, a codon supporting low levels of translation initiation by mitoribosomes *in vitro (37)*. We compared the *MT-ND5-dORF* AUG codon to other mtDNA AUG codons. In particular, we investigated those that are antisense to codons in known ORFs in an equivalent manner (i.e., those antisense to the second and third position of a codon and the first position of the subsequent codon). A large majority (82%) of these similarly-situated AUG codons had a smaller fraction of conservative changes across individuals (**Fig 1L**). Consequently, the *MT-ND5-dORF* start codon is retained in the human population, suggesting a possible functional role for *MT-ND5-dORF* translation.

### Balanced mitochondrial and cytosolic translation across cell types

Across bacteria and eukaryotes, subunits of most multiprotein complexes are synthesized proportionally to their final complex stoichiometry, with synthesis rates falling within a 2-fold range (*38, 39, 40*). Nuclear DNA-encoded mitochondrial complex subunits are a noted exception (*38*); consistent with this, we observed 2–32-fold differences in synthesis rates across subunits for each of the four dual-origin OXPHOS complexes (**Fig 2A,B**). Likewise, our mitoribosome profiling results showed that the mtDNA-encoded subunits were not synthesized proportionally to their complex stoichiometry, especially for Complex I (**Fig 2B**). Nevertheless, overall synthesis levels of each dual-origin complex follow the same pattern in the cytosol and mitochondria (**Fig S4, Table S2**). Indeed, average synthesis levels in the cytosol and mitochondria were highly correlated across complexes in all cell lines studied (**Fig 2C**). Mitochondrial and cytosolic synthesis levels were very highly correlated (Pearson’s r >0.95) in fibroblasts, myoblasts, myocytes, and HeLa S3 cells (**Fig 2C**) and only slightly less strongly correlated in HEK293T cells (Pearson’s r= 0.78) (**Fig 2C**). This translational balance was not solely due to RNA abundances, which were also positively correlated, albeit to a lesser extent (Pearson’s r= 0.55-0.79) (**Fig 2D**). Therefore, translation regulation promotes balanced synthesis of OXPHOS subunits across compartments in human cells.

**Figure 2.**
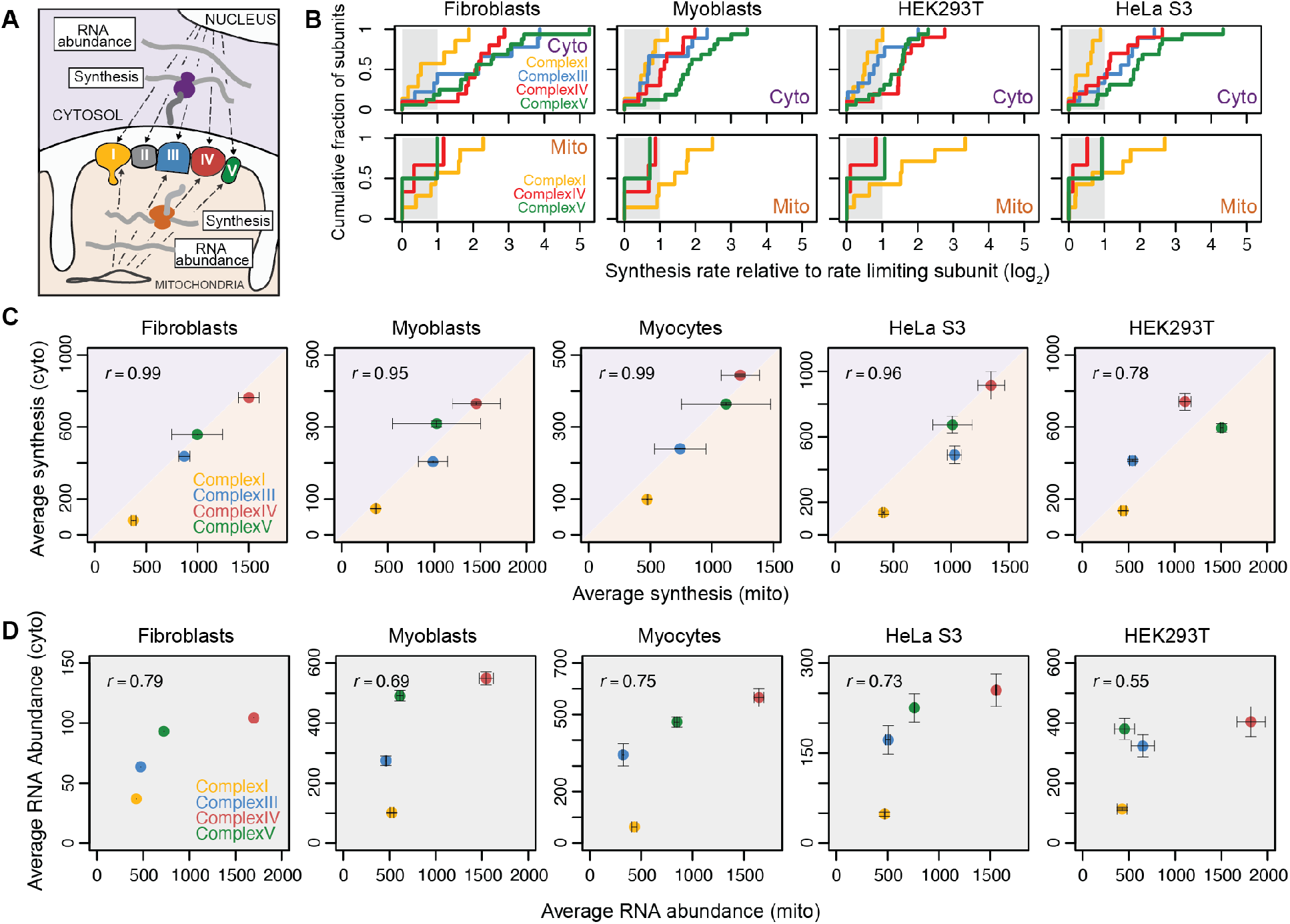
Balanced translatomes exist in five human cell lines. (**A**) Flow of genetic information to encode dual-origin OXPHOS complexes. (**B**) Cumulative distribution of relative synthesis rates for core nDNA-encoded subunits (top, see Figure S4A, Table S2 and Methods for details) and mtDNA-encoded subunits (bottom) of the dual-origin OXPHOS complexes. Log2-fold differences in synthesis rate compared to the rate-limiting (lowest synthesis rate) subunit of each complex are shown. Relative rates for cytosolic synthesis are measured as reads per kb per million mapped (RPKM), and for mitochondrial synthesis are measured as transcripts per ten thousand (tp10k, see Methods for details). Shaded region indicates 2-fold spread. (**C**) Average synthesis of mtDNA-encoded subunits (tp10k) compared to average synthesis of nDNA-encoded core subunits (RPKM) for each complex. Fibroblast and HeLa S3 cytoribosome profiling data are from (*55*) and (*54*), respectively. Myocytes are post-differentiation-induction day 2 myoblasts. Individual subunit synthesis for each replicate is shown in Figure S4A. Error bars show range across replicates. (**D**) Average RNA abundance of mtDNA-encoded subunits (tp10k) compared to average RNA abundance of nDNA-encoded core subunits (RPKM) for each complex. Subunits included are identical to those in (**C**). Error bars show range across replicates. All panels show Pearson correlation, r.

In yeast, cytosolic translation controls mitochondrial translation, directly facilitating balanced mitochondrial and cytosolic protein synthesis (*10, 11*). We asked whether cytosolic translation also controls mitochondrial translation in human cells in order to generate the correlated translatomes we observed. Inhibition of cytosolic translation for 30 minutes or 2 hours had no effect on relative or absolute mitochondrial protein synthesis (**Fig 3A, Fig S5A,D**). Similarly, inhibiting mitochondrial translation did not affect cytosolic translation in the short term (**Fig 3B, Fig S5B,C**). Thus, in contrast to the situation in yeast, we did not observe rapid communication from cytosolic to mitochondrial translation programs.

**Figure 3.**
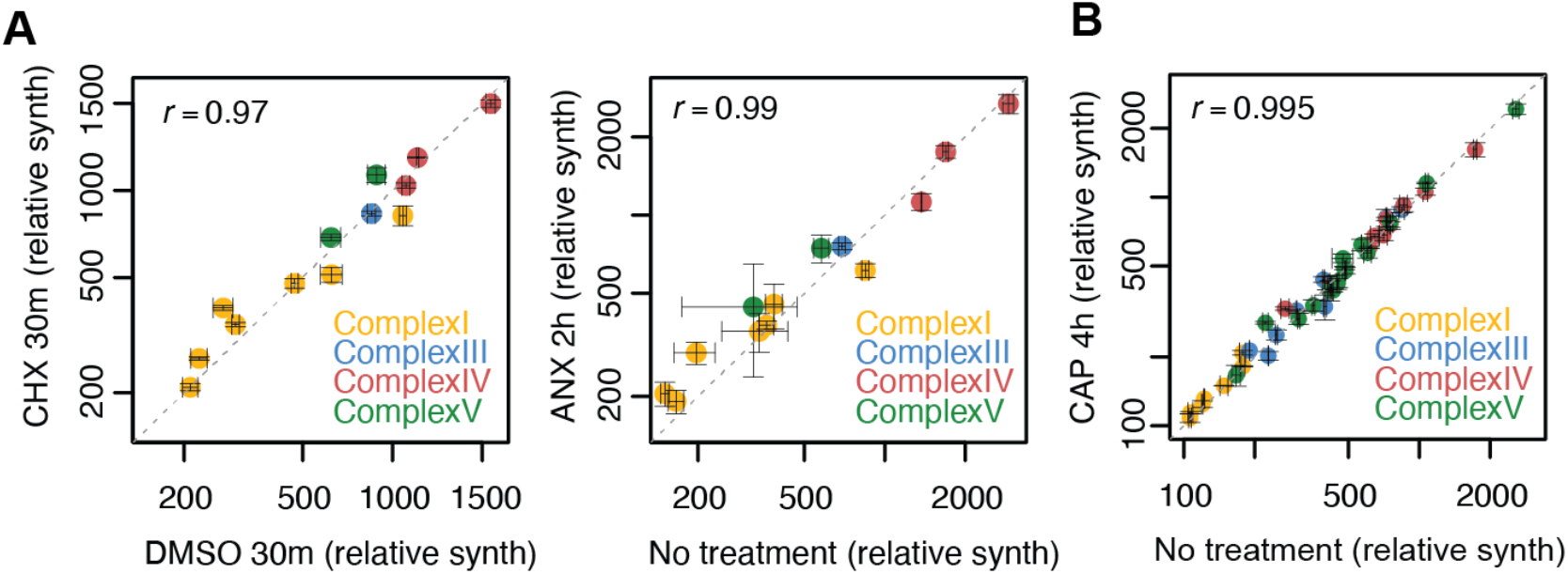
A lack of rapid communication between translation systems. (**A**) Comparison of relative synthesis rate values (tp10k) for mtDNA-encoded OXPHOS subunits with and without 30 minutes of cycloheximide (CHX) (100 µg/mL) or 2 hours of anisomycin (ANX) treatment (100 µg/mL) to inhibit cytosolic translation. (**B**) Comparison of relative synthesis rate values (RPKM) for cyto-translated OXPHOS subunits with and without 4 hours of chloramphenicol (CAP) treatment (100 µg/mL) to inhibit mitochondrial translation.

### LRPPRC is required for mitonuclear co-regulation

The RNA-binding protein LRPPRC regulates mitochondrial mRNA stability and interacts with the mitoribosome (*19–23*). Mutations in the *LRPPRC* gene lead to Leigh’s syndrome, a devastating neurometabolic disorder of childhood (*41*). To determine whether *LRPPRC* contributes to the coordination of the cytosolic and mitochondrial translatomes, we used CRISPR/Cas9 editing to generate single-cell clones in which expression of *LRPPRC* is ablated (**Fig S5E-G**), leading to extremely poor respiratory competence (∼10% of WT cells; **Fig S5H**). Mitochondrial and cytosolic protein synthesis became anticorrelated in the *LRPPRC*^KO^ cells (Pearson’s r = −0.89) (**Fig 4A**). RNA expression was also imbalanced, but not to the same degree (Pearson’s r = −0.21) **(Fig 4B)**. We also analyzed a cell line that overexpressed LRPPRC in the *LRPPRC*^KO^ background. The reconstituted cell line (*LRPPRC* rescue) restored the positively correlated mitochondrial and cytosolic translatomes observed in wild-type cells (**Fig 4A,B**). The OXPHOS malfunction in *LRPPRC*^KO^ cells may contribute to the imbalanced translatomes in these cells. Nevertheless, the established roles that LRPPRC plays in controlling mt-mRNA abundance and mitoribosome loading (*9-23*) suggest that *LRPPRC* directly aids in balancing mitochondrial translatomes and transcriptomes with their cytosolic counterparts.

**Figure 4.**
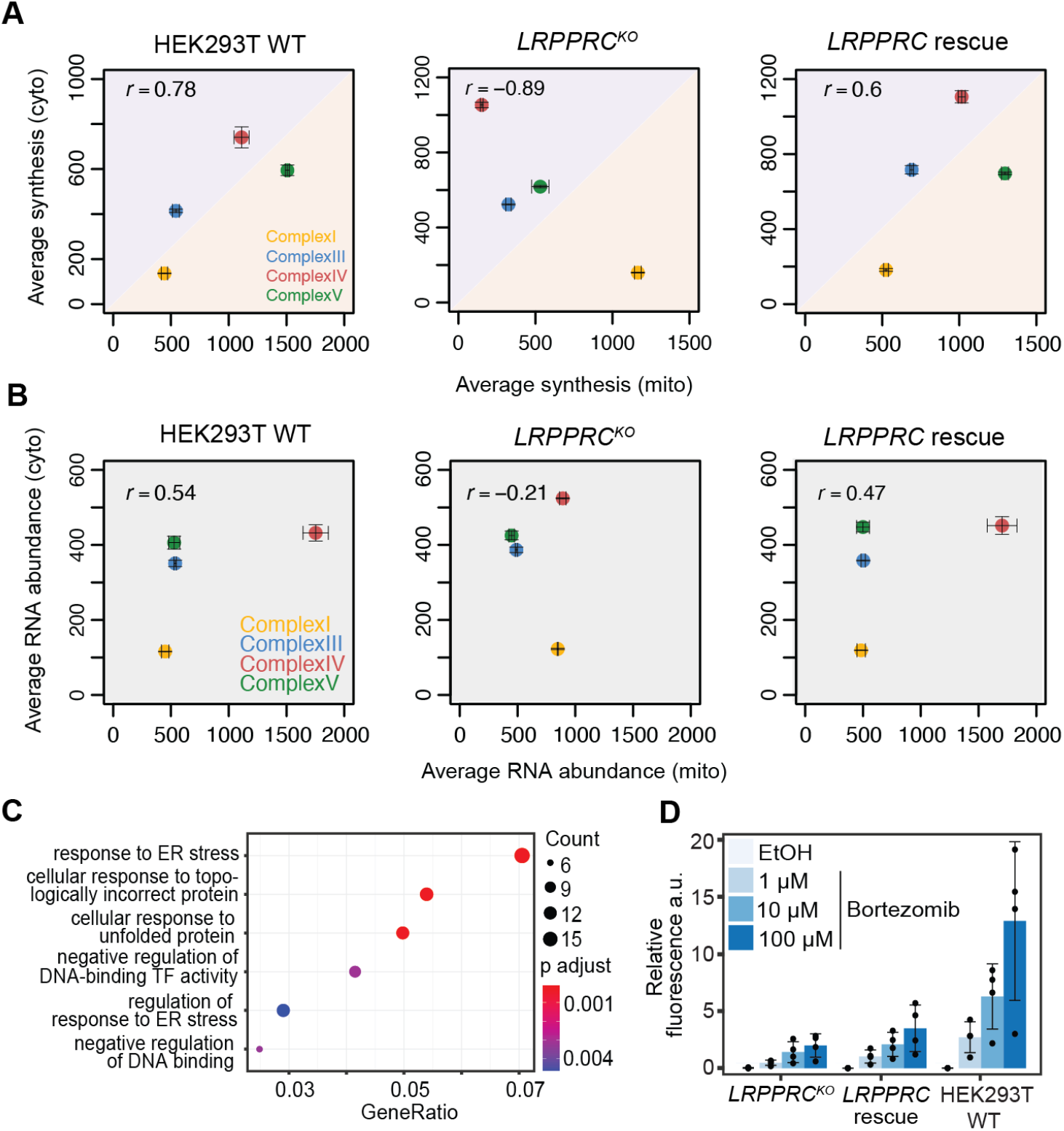
*LRPPRC* is necessary for mitonuclear balance. (**A**) Average synthesis of mtDNA-encoded subunits (tp10k) compared to average synthesis of nDNA-encoded core subunits (RPKM) for each complex in wild type HEK293T cells, *LRPPRC*^KO^ cells and the *LRPPRC*^KO^ cell line reconstituted for *LRPPRC* by its overexpression (*LRPPRC* rescue). (**B**) Average RNA abundance of mtDNA-encoded subunits (tp10k) compared to average RNA abundance of nDNA-encoded core subunits (RPKM) for each complex in *LRPPRC* cell lines. Error bars throughout show range across replicates, and all panels show Pearson correlation, r. (**C**) GO-term enrichment analysis of significantly (adjusted p-value < 0.05) differentially expressed genes that show a minimum two-fold increase in the *LRPPRC*^KO^ cells compared to the *LRPPRC* rescue cell line. Shown are the GO-terms after filtering for redundancy. p adjust = adjusted p-value. TF = transcription factor (**D**) Cell toxicity graph after 72 h bortezomib treatment. *LRPPRC* WT, *LRPPRC*^KO^ or *LRPPRC* rescue cells were treated with increasing amounts of bortezomib or the vehicle ethanol (EtOH). Cell death was measured via a fluorescence dye and living cells by indirect measurement of ATP levels via luminescence. Shown are the mean relative fluorescence arbitrary units (a.u.) normalized to the respective luminescence of 4 replicates. The range bars represent the standard deviation.

We asked whether a loss of balanced translatomes is associated with a loss of cellular proteostasis. Gene set enrichment analysis of *LRPPRC*^KO^ RNA-seq data revealed the up-regulation of genes involved in protein quality control, indicating a proteostasis strain exists in these cells (**Fig 4C, S5I, Tables S3, S4**). We observe the up-regulation of genes in the endoplasmic reticulum unfolded protein response pathway, consistent with its activation in yeast when nuclear DNA-encoded mitochondrial proteins are in excess (*42*) (**Fig 4C**). Moreover, these cells were less sensitive to bortezomib, a proteasome inhibitor, indicating that the induction of the protein quality control genes offered protection from further proteotoxicity (**Fig 4D**) (*43*). In sum, loss of *LRPPRC* disrupts mitonuclear translational balance and promotes the upregulation of proteostasis pathways, which in turn buffers cells from proteotoxicity.

## Discussion

A fundamental question in mitochondrial biology and cellular proteostasis is how the mitochondrial and nuclear genomes produce OXPHOS complexes in a coordinated fashion despite involving disparate gene expression systems operating in separate cellular compartments. In human cells, protein quality control pathways rapidly degrade orphaned OXPHOS proteins (*4, 9, 16*), and nDNA-encoded subunits of each OXPHOS complex are not synthesized in stoichiometric balance (*38*) (**Fig 2A,B**). These observations led to proposals that human cells may not need to coordinate cytosolic and mitochondrial protein synthesis (*4, 18*). In this study, we asked whether nuclear and mitochondrial OXPHOS genes are co-expressed in human cells. We found that mitochondrial and nuclear transcriptional programs produce OXPHOS RNA abundances that correspond modestly across complexes. Remarkably, translation regulation sharpens this correspondence so that the mitochondrial and cytosolic relative synthesis rates of the components of each complex are nearly perfectly correlated. Although we do not measure the absolute levels protein synthesis across compartments, the strong correlation across OXPHOS complexes suggests that the subunit flux from mitochondrial and cytosolic translation is matched for each respiratory complex, which presumably minimizes the risk of protein quality control overload. Indeed, loss of *LRPPRC* leads to mitonuclear imbalance and an upregulation of proteostasis pathways. These results are in line with observations in *C. elegans* and mouse cells, where disruption of mitonuclear balance leads to an upregulation of the mitochondrial unfolded protein response, which, in worms, extends lifespan (*6*).

In yeast, translation also plays a significant role in mitonuclear co-regulation of OXPHOS subunits (*10*). However, in that system, synchronous mitochondrial–cytosolic translation programs depend on cytosolic translation, indicating that direct communication between the translation systems enables the co-regulation. By contrast, in human cells, mitochondrial synthesis levels do not require ongoing cytosolic translation, at least on the time scale of hours. These differences are consistent with the paucity of mitochondrial translation regulators and UTRs that may represent a lack of direct control mechanisms for human mitochondrial translation. So how are mitochondrial and cytosolic translation fluxes maintained so that they correspond so precisely? We propose that on longer time scales, human mitochondrial translation adapts to the influx of nuclear DNA-encoded OXPHOS proteins (*44*), which requires the orchestrated effort of many mitochondrial gene expression regulators, including LRPPRC. How cross-compartment synthesis balance is detected and fine-tuned are important questions that should be addressed in future work. Finally, precisely balanced translatomes are likely a continual challenge for the cell to maintain across translation systems. Yet the consequence of imbalance is high -- proteostasis collapse. Thus, we anticipate that the tight reciprocity of mitochondrial and cytosolic translatomes represents a key vulnerability in cellular proteostasis and mitochondrial function.

## Acknowledgements

We thank Dan Davidi for analysis help; Jake Bridgers, Karine Choquet, Stefan Isaacs and Dan Davidi for manuscript comments; Chris Patil for manuscript editing.

## Author contributions

L.S.C and I.S. and M.C. conceived the study; I.S., E.M. and K.G.H. collected ribosome profiling and RNA-seq data; K.G.H. performed cell survival assays, I.S. and K.G.H. analyzed the data; M.C. created software; I.S., M.C., E.M. and K.G.H. made the figures; J.C.M. and A.B. created and characterized the CRISPR/Cas9 *LRPPRC*^*KO*^ cell lines; L.S.C, I.S., M.C. and E.M. wrote the paper. All authors edited the manuscript.

## Data and materials availability

Raw and processed sequencing data were deposited in the GEO database under the accession number GSE173283. Scripts and instructions for human mitoribosome profiling data analyses are available at https://github.com/churchmanlab/human-mitoribosome-profiling.

## Supplementary Materials

Materials and Methods

Figs. S1 to S5

Tables S1 to S4

## References and Notes

1. S. Rath, R. Sharma, R. Gupta, T. Ast, C. Chan, T. J. Durham, R. P. Goodman, Z. Grabarek, M. E. Haas, W. H. W. Hung, P. R. Joshi, A. A. Jourdain, S. H. Kim, A. V. Kotrys, S. S. Lam, J. G. McCoy, J. D. Meisel, M. Miranda, A. Panda, A. Patgiri, R. Rogers, S. Sadre, H. Shah, O. S. Skinner, T.-L. To, M. A. Walker, H. Wang, P. S. Ward, J. Wengrod, C.-C. Yuan, S. E. Calvo, V. K. Mootha, MitoCarta3.0: an updated mitochondrial proteome now with sub-organelle localization and pathway annotations. Nucleic Acids Res. 49, D1541–D1547 (2021).

2. M. T. Ryan, N. J. Hoogenraad, Mitochondrial-nuclear communications. Annu. Rev. Biochem. 76, 701–722 (2007).

3. A. Mottis, S. Herzig, J. Auwerx, Mitocellular communication: Shaping health and disease. Science. 366, 827–832 (2019).

4. R. S. Isaac, E. McShane, L. S. Churchman, The Multiple Levels of Mitonuclear Coregulation. Annu. Rev. Genet. 52, 511–533 (2018).

5. A. P. Gomes, N. L. Price, A. J. Y. Ling, J. J. Moslehi, M. K. Montgomery, L. Rajman, J. P. White, J. S. Teodoro, C. D. Wrann, B. P. Hubbard, E. M. Mercken, C. M. Palmeira, R. de Cabo, A. P. Rolo, N. Turner, E. L. Bell, D. A. Sinclair, Declining NAD+ Induces a Pseudohypoxic State Disrupting Nuclear-Mitochondrial Communication during Aging. Cell. 155, 1624–1638 (2013).

6. R. H. Houtkooper, L. Mouchiroud, D. Ryu, N. Moullan, E. Katsyuba, G. Knott, R. W. Williams, J. Auwerx, Mitonuclear protein imbalance as a conserved longevity mechanism. Nature. 497, 451–457 (2013).

7. G. C. Kujoth, A. Hiona, T. D. Pugh, S. Someya, K. Panzer, S. E. Wohlgemuth, T. Hofer, A. Y. Seo, R. Sullivan, W. A. Jobling, J. D. Morrow, H. Van Remmen, J. M. Sedivy, T. Yamasoba, M. Tanokura, R. Weindruch, C. Leeuwenburgh, T. A. Prolla, Mitochondrial DNA mutations, oxidative stress, and apoptosis in mammalian aging. Science. 309, 481–484 (2005).

8. O. Khalimonchuk, A. Bird, D. R. Winge, Evidence for a Pro-oxidant Intermediate in the Assembly of Cytochrome Oxidase. J. Biol. Chem. 282, 17442–17449 (2007).

9. J. Song, J. M. Herrmann, T. Becker, Quality control of the mitochondrial proteome. Nat. Rev. Mol. Cell Biol. 22, 54–70 (2021).

10. M. T. Couvillion, I. C. Soto, G. Shipkovenska, L. S. Churchman, Synchronized mitochondrial and cytosolic translation programs. Nature. 533, 499–503 (2016).

11. J. M. Herrmann, M. W. Woellhaf, N. Bonnefoy, Control of protein synthesis in yeast mitochondria: the concept of translational activators. Biochim. Biophys. Acta. 1833, 286–294 (2013).

12. M. Ott, A. Amunts, A. Brown, Organization and Regulation of Mitochondrial Protein Synthesis. Annu. Rev. Biochem. 85, 77–101 (2016).

13. M. C. Costanzo, T. D. Fox, Product of Saccharomyces cerevisiae nuclear gene PET494 activates translation of a specific mitochondrial mRNA. Mol. Cell. Biol. 6, 3694–3703 (1986).

14. A. L. Moyer, K. R. Wagner, Mammalian Mss51 is a skeletal muscle-specific gene modulating cellular metabolism. J Neuromuscul Dis. 2, 371–385 (2015).

15. W. Weraarpachai, H. Antonicka, F. Sasarman, J. Seeger, B. Schrank, J. E. Kolesar, H. Lochmüller, M. Chevrette, B. A. Kaufman, R. Horvath, E. A. Shoubridge, Mutation in TACO1, encoding a translational activator of COX I, results in cytochrome c oxidase deficiency and late-onset Leigh syndrome. Nat. Genet. 41, 833–837 (2009).

16. D. F. Bogenhagen, J. D. Haley, Pulse-chase SILAC-based analyses reveal selective oversynthesis and rapid turnover of mitochondrial protein components of respiratory complexes. J. Biol. Chem. 295, 2544–2554 (2020).

17. P. M. Quirós, A. Mottis, J. Auwerx, Mitonuclear communication in homeostasis and stress. Nat. Rev. Mol. Cell Biol. 17, 213–226 (2016).

18. E. Kummer, N. Ban, Mechanisms and regulation of protein synthesis in mitochondria. Nat. Rev. Mol. Cell Biol. 22, 307–325 (2021).

19. S. Aibara, V. Singh, A. Modelska, A. Amunts, Structural basis of mitochondrial translation. Elife. 9 (2020), doi:10.7554/eLife.58362.

20. M. Lagouge, A. Mourier, H. J. Lee, H. Spåhr, T. Wai, C. Kukat, E. Silva Ramos, E. Motori, J. D. Busch, S. Siira, German Mouse Clinic Consortium, E. Kremmer, A. Filipovska, N.-G. Larsson, SLIRP Regulates the Rate of Mitochondrial Protein Synthesis and Protects LRPPRC from Degradation. PLoS Genet. 11, e1005423 (2015).

21. T. Chujo, T. Ohira, Y. Sakaguchi, N. Goshima, N. Nomura, A. Nagao, T. Suzuki, LRPPRC/SLIRP suppresses PNPase-mediated mRNA decay and promotes polyadenylation in human mitochondria. Nucleic Acids Res. 40, 8033–8047 (2012).

22. F. Sasarman, C. Brunel-Guitton, H. Antonicka, T. Wai, E. A. Shoubridge, LSFC Consortium, LRPPRC and SLIRP interact in a ribonucleoprotein complex that regulates posttranscriptional gene expression in mitochondria. Mol. Biol. Cell. 21, 1315–1323 (2010).

23. B. Ruzzenente, M. D. Metodiev, A. Wredenberg, A. Bratic, C. B. Park, Y. Cámara, D. Milenkovic, V. Zickermann, R. Wibom, K. Hultenby, H. Erdjument-Bromage, P. Tempst, U. Brandt, J. B. Stewart, C. M. Gustafsson, N.-G. Larsson, LRPPRC is necessary for polyadenylation and coordination of translation of mitochondrial mRNAs. EMBO J. 31, 443– 456 (2012).

24. Y. Itoh, J. Andréll, A. Choi, U. Richter, P. Maiti, R. B. Best, A. Barrientos, B. J. Battersby, A. Amunts, Mechanism of membrane-tethered mitochondrial protein synthesis. Science. 371, 846–849 (2021).

25. N. T. Ingolia, G. A. Brar, S. Rouskin, A. M. McGeachy, J. S. Weissman, Curr. Protoc. Mol. Biol., Unit 4.18 (2013).

26. D. Ojala, G. Attardi, Expression of the mitochondrial genome in HeLa cells. X. Properties of mitochondrial polysomes. J. Mol. Biol. 65, 273–289 (1972).

27. R. Englmeier, S. Pfeffer, F. Förster, Structure of the Human Mitochondrial Ribosome Studied In Situ by Cryoelectron Tomography. Structure. 25, 1574–1581.e2 (2017).

28. U. Richter, M. E. Evans, W. C. Clark, P. Marttinen, E. A. Shoubridge, A. Suomalainen, A. Wredenberg, A. Wedell, T. Pan, B. J. Battersby, RNA modification landscape of the human mitochondrial tRNALys regulates protein synthesis. Nat. Commun. 9, 3966 (2018).

29. M. Lu, L. Zhou, W. C. Stanley, M. E. Cabrera, G. M. Saidel, X. Yu, Role of the malate-aspartate shuttle on the metabolic response to myocardial ischemia. J. Theor. Biol. 254, 466– 475 (2008).

30. F. Gao, M. Wesolowska, R. Agami, K. Rooijers, F. Loayza-Puch, C. Lawless, R. N. Lightowlers, Z. M. A. Chrzanowska-Lightowlers, Using mitoribosomal profiling to investigate human mitochondrial translation. Wellcome Open Res. 2, 116 (2017).

31. B. E. Christian, L. L. Spremulli, Preferential selection of the 5’-terminal start codon on leaderless mRNAs by mammalian mitochondrial ribosomes. J. Biol. Chem. 285, 28379– 28386 (2010).

32. D. Ojala, J. Montoya, G. Attardi, tRNA punctuation model of RNA processing in human mitochondria. Nature. 290, 470–474 (1981).

33. A. R. Wolf, V. K. Mootha, Functional genomic analysis of human mitochondrial RNA processing. Cell Rep. 7, 918–931 (2014).

34. J. R. Wangen, R. Green, Stop codon context influences genome-wide stimulation of termination codon readthrough by aminoglycosides. Elife. 9 (2020), doi:10.7554/eLife.52611.

35. B. Miller, S.-J. Kim, H. Kumagai, H. H. Mehta, W. Xiang, J. Liu, K. Yen, P. Cohen, Peptides derived from small mitochondrial open reading frames: Genomic, biological, and therapeutic implications. Exp. Cell Res. 393, 112056 (2020).

36. A. Bolze, F. Mendez, S. White, F. Tanudjaja, M. Isaksson, R. Jiang, A. D. Rossi, E. T. Cirulli, M. Rashkin, W. J. Metcalf, J. J. Grzymski, W. Lee, J. T. Lu, N. L. Washington, A catalog of homoplasmic and heteroplasmic mitochondrial DNA variants in humans. bioRxiv (2020), p. 798264.

37. M. Lee, N. Matsunaga, S. Akabane, I. Yasuda, T. Ueda, N. Takeuchi-Tomita, Reconstitution of mammalian mitochondrial translation system capable of correct initiation and long polypeptide synthesis from leaderless mRNA. Nucleic Acids Res. 49, 371–382 (2021).

38. J. C. Taggart, G.-W. Li, Production of Protein-Complex Components Is Stoichiometric and Lacks General Feedback Regulation in Eukaryotes. Cell Syst. 7, 580–589.e4 (2018).

39. G.-W. Li, D. Burkhardt, C. Gross, J. S. Weissman, Quantifying absolute protein synthesis rates reveals principles underlying allocation of cellular resources. Cell. 157, 624–635 (2014).

40. J. C. Taggart, H. Zauber, M. Selbach, G.-W. Li, E. McShane, Keeping the Proportions of Protein Complex Components in Check. Cell Syst. 10, 125–132 (2020).

41. V. K. Mootha, P. Lepage, K. Miller, J. Bunkenborg, M. Reich, M. Hjerrild, T. Delmonte, A. Villeneuve, R. Sladek, F. Xu, G. A. Mitchell, C. Morin, M. Mann, T. J. Hudson, B. Robinson, J. D. Rioux, E. S. Lander, Identification of a gene causing human cytochrome c oxidase deficiency by integrative genomics. Proc. Natl. Acad. Sci. U. S. A. 100, 605–610 (2003).

42. K. Knöringer, C. Groh, L. Krämer, K. C. Stein, K. G. Hansen, J. M. Herrmann, J. Frydman, F. Boos, The unfolded protein response of the endoplasmic reticulum supports mitochondrial biogenesis by buffering non-imported proteins. bioRxiv (2021), p. 2021.05.19.444788.

43. M. Groll, C. R. Berkers, H. L. Ploegh, H. Ovaa, Crystal structure of the boronic acid-based proteasome inhibitor bortezomib in complex with the yeast 20S proteasome. Structure. 14, 451–456 (2006).

44. R. Richter-Dennerlein, S. Oeljeklaus, I. Lorenzi, C. Ronsör, B. Bareth, A. B. Schendzielorz, C. Wang, B. Warscheid, P. Rehling, S. Dennerlein, Mitochondrial Protein Synthesis Adapts to Influx of Nuclear-Encoded Protein. Cell. 167, 471–483.e10 (2016).

45. N. T. Ingolia, G. A. Brar, S. Rouskin, A. M. McGeachy, J. S. Weissman, The ribosome profiling strategy for monitoring translation in vivo by deep sequencing of ribosome-protected mRNA fragments. Nat. Protoc. 7, 1534–1550 (2012).

46. R. Shalgi, J. A. Hurt, I. Krykbaeva, M. Taipale, S. Lindquist, C. B. Burge, Widespread regulation of translation by elongation pausing in heat shock. Mol. Cell. 49, 439–452 (2013).

47. M. Martin, Cutadapt removes adapter sequences from high-throughput sequencing reads. EMBnet.journal. 17, 10–12 (2011).

48. B. Langmead, C. Trapnell, M. Pop, S. L. Salzberg, Ultrafast and memory-efficient alignment of short DNA sequences to the human genome. Genome Biol. 10, R25 (2009).

49. A. Dobin, C. A. Davis, F. Schlesinger, J. Drenkow, C. Zaleski, S. Jha, P. Batut, M. Chaisson, T. R. Gingeras, STAR: ultrafast universal RNA-seq aligner. Bioinformatics. 29, 15–21 (2013).

50. K. Rooijers, F. Loayza-Puch, L. G. Nijtmans, R. Agami, Ribosome profiling reveals features of normal and disease-associated mitochondrial translation. Nat. Commun. 4, 1–8 (2013).

51. S. Gopalakrishna, S. F. Pearce, A. M. Dinan, F. A. Schober, M. Cipullo, H. Spåhr, A. Khawaja, C. Maffezzini, C. Freyer, A. Wredenberg, I. Atanassov, A. E. Firth, J. Rorbach, C6orf203 is an RNA-binding protein involved in mitochondrial protein synthesis. Nucleic Acids Res. 47, 9386–9399 (2019).

52. S. F. Pearce, J. Rorbach, L. Van Haute, A. R. D’Souza, P. Rebelo-Guiomar, C. A. Powell, I. Brierley, A. E. Firth, M. Minczuk, Maturation of selected human mitochondrial tRNAs requires deadenylation. Elife. 6 (2017), doi:10.7554/eLife.27596.

53. S. H.-J. Li, M. Nofal, L. R. Parsons, J. D. Rabinowitz, Z. Gitai, Monitoring mammalian mitochondrial translation with MitoRiboSeq. Nat. Protoc. (2021), doi:10.1038/s41596-021-00517-1.

54. C. C.-C. Wu, B. Zinshteyn, K. A. Wehner, R. Green, High-Resolution Ribosome Profiling Defines Discrete Ribosome Elongation States and Translational Regulation during Cellular Stress. Mol. Cell. 73, 959–970.e5 (2019).

55. O. Tirosh, Y. Cohen, A. Shitrit, O. Shani, V. T. K. Le-Trilling, M. Trilling, G. Friedlander, M. Tanenbaum, N. Stern-Ginossar, The Transcription and Translation Landscapes during Human Cytomegalovirus Infection Reveal Novel Host-Pathogen Interactions. PLoS Pathog. 11, e1005288 (2015).

56. Y. Liao, J. Wang, E. J. Jaehnig, Z. Shi, B. Zhang, WebGestalt 2019: gene set analysis toolkit with revamped UIs and APIs. Nucleic Acids Res. 47, W199–W205 (2019).

57. P. Jesina, M. Tesarová, D. Fornůsková, A. Vojtísková, P. Pecina, V. Kaplanová, H. Hansíková, J. Zeman, J. Houstek, Diminished synthesis of subunit a (ATP6) and altered function of ATP synthase and cytochrome c oxidase due to the mtDNA 2 bp microdeletion of TA at positions 9205 and 9206. Biochem. J. 383, 561–571 (2004).

58. M. I. Love, W. Huber, S. Anders, Moderated estimation of fold change and dispersion for RNA-seq data with DESeq2. Genome Biol. 15, 550 (2014).

59. G. Yu, L.-G. Wang, Y. Han, Q.-Y. He, clusterProfiler: an R package for comparing biological themes among gene clusters. OMICS. 16, 284–287 (2012).

