## Supplemental materials for "Balanced mitochondrial and cytosolic translatomes underlie the biogenesis of human respiratory complexes"

**This PDF file includes:**

Materials and Methods
Figs. S1 to S5
Legends for Tables S1 to S4

**Other Supplementary Materials for this manuscript include the following:**

Tables S1 to S4 [TableS1.xlsx, TableS2.xlsx, TableS3.xlsx, TableS4.xlsx]

#### Materials and Methods

##### Cell cultures

Dermal human fibroblasts, HeLa S3, Mouse NIH 3T3 and HEK293T were grown in DMEM (Thermo, CAT 11965092), supplemented with 10% FBS (Thermo Fisher, CAT A3160402). Human myoblasts and myocytes were grown in Human Skeletal Muscle Basal Media Kit (Cell Applications, CAT 151K-500). Unless otherwise indicated, all cell cultures were grown in 15 cm dishes at 37 $^0$C, 5% $CO_2$, in the absence of antibiotics to 75-80% confluency. Dermal human fibroblasts and myoblasts were anonymous healthy control samples kindly provided by Dr. Brendan Battersby (Institute of Biotechnology, University of Helsinki.)

##### Myoblast differentiation to Myocytes

Human myoblasts were cultured in Human Skeletal Muscle Basal Media Kit (Cell Applications, CAT 151K-500) to 80% confluency. Differentiation was induced by shifting cell cultures to rich medium (DMEM, Thermo Fisher Scientific, CAT 11965092) supplemented with 2% horse serum (Thermo Fisher Scientific, CAT 26050070) and 0.4 μg/mL dexamethasone (Sigma, CAT D4902). Myocytes were harvested two days post-differentiation-induction.

##### Drug treatments

Cells exposed to ribosome inhibitors were grown to 70% confluency, as explained above, before addition of the drug. Growth medium was replaced by a medium containing 100 μg/mL of the inhibitor [cycloheximide (Sigma, CAT C769), chloramphenicol (Sigma, CAT C3175), or anisomycin (Sigma, CAT A9789)], and 100 μg/mL of uridine (Sigma, CAT U3750). Treatments were done under the same incubation conditions as growth, for the indicated times.

##### Generation of mutant cell lines

To create the *LRPPRC* knockout, we first obtained a pool of HEK293T cells that had been transfected with the guide RNA 5'-GAGGACUACUGAGCCCAGCC-3' targeting exon 2, from Synthego (Menlo Park, CA). This pool was then plated into 96 well plates to screen for individual clones and these clones were screened for presence of LRPPRC by western blot (Santa Cruz, sc-166178). Clones that showed absence of LRPPRC by western blot were then genotyped by subcloning the edited section into the pCR 2.1-TOPO TA vector (Thermo-Fisher CAT:451641). Clones were then picked and the plasmid was sequenced by Sanger sequencing (GeneWiz). To create the reconstituted cell line, a Myc-DDK tagged *LRPPRC* ORF plasmid was obtained from OriGene (CAT: RC216747). This ORF was then sub cloned into a hygromycin resistance-containing pCMV6 entry vector (OriGene, CAT PS100024). $LRPPRC^{KO}$ cells were then transfected with 1.5 µg of plasmid using endofectin (GeneCopoeia, CAT EF014) as the transfection agent. They were then selected for two weeks on 200 µg/mL hygromycin (Invitrogen, CAT 10687010) and LRPPRC levels were checked by western blot. For experiments conducted with the $LRPPRC^{KO}$ , and control cell lines (WT HEK293T and reconstituted $LRPPRC^{KO}$ +WT rescue), we cultured cells in DMEM (Thermo Fisher Scientific, CAT 11965092), supplemented with 10% FBS (Thermo Fisher, CAT A3160402), 100 µg/mL of uridine (Sigma, CAT U3750), 3 mM sodium formate (Sigma CAT 247596), 1 mM sodium pyruvate (Thermo Fisher Scientific, CAT 11360070) and 1 mM GlutaMAX (Thermo Fisher Scientific, CAT 35050061).

Polarographic measurements in mutant cell lines

Endogenous cell respiration was measured polarographically at $37^0$C using a Clark-type electrode from Hansatech Instruments (Norfolk, United Kingdom). Briefly, trypsinized cells were washed with permeabilized-cell respiration buffer (PRB) containing 0.3 M mannitol, 10 mM KCl, 5 mM $MgCl_2$, 0.5 mM EDTA, 0.5 mM EGTA, 1 mg/ml BSA and 10 mM $KH_3PO_4$(pH 7.4). The cells were resuspended at approximately $2x10^6$ cells/ml in 0.5 mL of the same buffer air-equilibrated at

$37^0C$. The cell suspension was immediately placed into the polarographic chamber to measure endogenous respiration. Subsequently, complex IV activity was inhibited using 0.8 mM KCN to assess the mitochondrial specificity of the oxygen consumption measured. Values were normalized by total cell number.

Cell toxicity assay

To test the toxicity of Bortezomib (Cayman Chemical Company, CAT 10008822), WT HEK293T, *LRPPRC*$^{KO}$ HEK293T or *LRPPRC*$^{KO}$ rescue cells were grown in a black-walled 96 well plate (Corning Incorporated costar®, CAT CLS3603-48EA). The same growth media was used as described in the mutant generation section. 5000 cells per well were seeded in a 50 µl volume and left to adhere for 1 to 2 h. The adherence of the cells was checked by visual inspection by light microscopy. Media supplemented with increasing amounts of the protease inhibitor Bortezomib (final concentrations 1 µM, 10 µM and 100 µM) were added to the well. The vehicle ethanol (EtOH) was used as a control (final concentration 1% (v/v)). Cells were grown for 72 hours. The toxicity of the drug was measured using the CellTox™ Green Cytotoxicity Assay from Promega (CAT G8742) as described in the instructions. In brief, the dye was mixed with the assay buffer added to a final 1x concentration and the plate was transferred into a plate reader (Tecan Infinite 200 Pro). To allow for mixing, the plate was incubated with shaking for 60 s at an amplitude of 1 mm, followed by 14 min wait time, and another 60 s of shaking. After 10 s of settling, the fluorescence was measured using a 485 nm emission and 535 nm excitation filter set. Since the cell toxicity measures the amount of accumulated dead cells, the cell number needs to be determined. To this end, the CellTiter-Glo® Luminescent Cell Viability Assay from Promega (CAT G7571) was used to measure ATP levels for an estimate of living cells. Here, the 2x reagent was added to the before mix and the plate was transferred into the plate reader. To mix the reagent with the media, the plate was incubated with shaking for 70 to 120 s at 1 mm amplitude followed by a 10 min incubation without shaking. Afterwards, luminescence was measured. For analysis, first the background was subtracted from raw fluorescence or raw luminescence values. The fluorescence of each well was normalized to the luminescence measured of the respective well. Each biological replicate (n=4) includes two to three technical replicates. Analysis was performed in R.

General nucleic acid and protein methods

Northern blots were performed to assess mitoribosome footprint size. We isolated RNA from mitomonosome-enriched sucrose gradient fractions by extracting with 1:1 volume of acid-phenol/chloroform (Thermo-Fisher Scientific). Purified RNA was separated on a 15% polyacrylamide TBE-urea gel. After transfer to a nylon membrane (Hybond N+), blots were probed at room temperature with $^{32}$P-dATP-internally labelled (Perkin Elmer) random hexamer-primed DNA fragments synthesized from 400–500-bp PCR-generated templates for *MT-CO2*. Blots were exposed to a Storage Phosphor Screen and visualized using a Typhoon instrument.

Proteins were resolved before western blotting on NuPAGE Novex Bis-Tris gels (Thermo Fisher Scientific). After transfer to a nitrocellulose membrane, blots were probed for primary antibodies against mitoribosome subunits MRPL12 (Proteintech, 14795-1-AP) and MRPS18-B (Proteintech, 16139-1-AP). Other antibodies used: COI (ABCAM, ab14705), HSP60 (Cell Signaling, 4870S) and LRPPRC (Proteintech, 21175-1-AP). Fluorescently labelled secondary antibodies (IRDye, LI-COR) were detected using a LI-COR Odyssey instrument.

*In vivo* labeling of translation products

Cells were grown in FBS-supplemented complete DMEM, as explained in the cell culture section above, to 80% confluency. To equilibrate cells to the labeling conditions, growth medium was replaced with labeling medium (DMEM depleted of cysteine, Thermo Fisher Scientific CAT

21013-024) 30 minutes before beginning of labeling. After 30 minutes of incubation at 37 $^{0}$C, 5 % $CO_2$, the corresponding cytosolic translation inhibitor was added to the plates to a final concentration of 100 μg/ml. Treatment was performed for the indicated times. After treatment completion, 200 μCi/mL of EasyTag labeling mix (35S-cysteine/35S-methionine, Perkin Elmer CAT NEG772007MC) was added to each plate. Labeling was followed for the indicated times. To collect, cells were rinsed twice with cold PBS 1X pH:7.4, in 1 mL of 1X PBS pH:7.4, scraped using a cell lifter and pelleted by centrifugation at 1,500 g for 10 min at 4ºC. To prepare samples for electrophoresis, cell pellets were extracted with 1X gel loading buffer (95 mM Tris–HCl, pH:6.8, 7.5% glycerol, 2% SDS, 0.5 mg/ml bromophenol blue, 50 mM DTT). Samples were separated in a 17.5% acrylamide gel (20 cm). After transferring to nitrocellulose membrane, signals were collected by a phosphor imager screen and visualized using a Typhoon instrument.

Mitoribosome profiling

Cell cultures of biological replicates were grown independently in 15 cm dishes to 70-80% confluence. Medium was aspirated and cells were quickly rinsed once with ice-cold 1X PBS pH 7.4. Lysis was performed on ice in a glass homogenizer with a loose setting. We used 50 ml of lysis buffer for $10^6$ cells. Example: 500 μL to lyse a 15 cm dish of human fibroblasts and 1000 μL to lyse a 15 cm dish of HeLa S3 cells. Lysis buffer composition was carefully designed to preserve the integrity on the mitoribosome. We used lauryl maltoside at 0.25% to fully solubilize mitochondrial membranes, while preserving their integrity, in combination with 50 mM $NH_4Cl$, 20 mM $MgCl_2$, 0.5 mM DTT, 10 mM Tris, pH 7.5 and 1X EDTA-free Protease inhibitor cocktail (Roche). To normalize for global changes in mitochondrial translation, we introduced a mouse spike-in control. Before nuclease digestion, mouse (NIH 3T3) lysate was mixed in with the human at 95%:5% (human:mouse) $OD_{260}$ equivalent volumes. Mitoribosome footprints were generated using 8 Units/mL of RNase If (NEB), digesting 450 μL of whole cell lysate at room temperature for 30 minutes, without rotation. Digestion was stopped by adding 80 Units of SUPERaseIn (Thermo Fisher Scientific). To isolate mitoribosomes, lysates were clarified at 10,000 RPM for 5 minutes at $4^0$C, loaded on 10-50% linear sucrose gradients and centrifuged in a Beckman ultra-centrifuge at 40,000 RPM for three hours at $4^0$C using a SW 41 Ti rotor. Gradients were mixed and fractionated using a BioComp instrument. The mitomonosome fraction was identified using western blots for MRPL12 and MRPS-18 (see protein methods above). Mitoribosome footprints were recovered by 1:1 volume phenol/chloroform extraction of the monosome fraction. RNA was separated in 15% polyacrylamide TBE-urea gels, excising sizes between 28-40 nucleotides. Libraries were prepared as described in (*45*), with a few modifications: (1) We did not rRNA deplete and (2) We introduced 6 or 10 randomized nucleotides to the linker ligation at 3' (/5rApp/NNNNNNNNNNCTGTAGGCACCATCAAT/3ddC/ or /5rApp/NNNNNNCTGTAGGCACCATCAAT/3ddC/) and in many samples also at the reverse transcription step at 5'(/5Phos/NNNNGATCGTCGGACTGTAGAACTCTGAACCTGTC/iSp18/CACTCA/iSp18/CAAGCAGAAGACGGCATACGAGATATTGATGGTGCCTACAG). Sequencing was performed in an Illumina based Next-seq 500 instrument at the Bauer sequencing facility (at Harvard University).

<u>Mitoribosome immunoprecipitation</u>

The mitoribosome was immunoprecipitated from the monosome-enriched sucrose fraction using a MPL12 antibody (Proteintech) conjugated to a slurry of DynaBeads protein A from Thermo Fisher Scientific. Mitoribosomes were recovered by eluting beads with mitoribosome profiling lysis buffer without lauryl maltoside but containing 0.1% SDS. Eluted mitoribosome footprints were extracted with 1:1 volume of phenol/chloroform. Libraries were prepared and sequenced as described in the mitoribosome profiling section above.

Cytoribosome profiling

Cell culture growth and lysis were performed as for mitoribosome profiling (see mitoribosome profiling section above). We used lysis and buffer conditions described in (*46*) for polysome profiling. Cytoribosome footprints were generated using $8x10^{-4}$ Units/mL of RNase I (Epicentre), digesting 450 µL of lysate at room temperature for 30 minutes, without rotation. Digestion was stopped by adding 80 Units of SUPERaseIn (Thermo Fisher). Isolation of the cytomonosome was performed using the centrifugation and fractionation conditions described for mitoribosome profiling but limiting the centrifugation time to two and a half hours. The cytomonosome fraction was detected by their UV absorbance at 254 nm. Cytoribosome footprints recovery and gel size purification were performed as in mitoribosome profiling but restricting the excision size to 27-33 nucleotides. Library preparation and sequencing techniques were identical to those used for mitoribosome profiling.

RNA-seq

Some of the lysates prepared for mitoribosome profiling were reserved for RNA-seq. Lysates were further treated with 90 µg/mL of Proteinase K and 0.5% SDS at 42 $^{0}$C for 20 minutes. Total RNA was extracted using 1:1 volume of phenol/chloroform. Purified RNA was DNase digested at 37 $^{0}$C for 30 minutes with 3 Units of RQ1 RNase-free DNase (Promega). We then rRNA-depleted 2 µg of RNA using the RiboMinus Eukaryotic kit v2 from Thermo Fisher (CAT: A15026). RNA was fragmented by alkaline hydrolysis in 5 mM $Na_2CO_3$, 45 mM $NaHCO_3$, 1 mM EDTA, pH 9.3 for 25 min at 95 °C. RNA was resolved in 15% TBE-Urea gels and size purified including 30–70-nucleotide fragments. Sequencing libraries were prepared and sequenced as above, except for HeLa S3 samples, which were size selected and prepared using the Illumina TruSeq kit.

Mitoribosome profiling data analysis

Reads were trimmed to remove ligated 3' linker (CTGTAGGCACCATCAAT) with Cutadapt (*47*). Reads without linker were discarded and the unique molecular identifier (UMI) was then extracted from remaining reads using a custom script, but kept associated with each read entry. UMIs consisted of either 6 or 10 random nucleotides at the 3' end ligated as part of the 3' linker, and either 0 or 4 random nucleotides at the 5' end ligated during circularization and originating from the RT primer. Trimmed reads were filtered after alignment using bowtie1 (*48*) to human rRNA (allowing one internal and one 5' mismatch) and human tRNA (allowing one internal, one 5', and three 3' mismatches). Remaining reads were aligned to the GRCh38 human reference genome merged with the M17 mouse reference genome from GENCODE using STAR 2.7.3 (*49*) with parameters --outFilterMismatchNoverReadLmax 0.07 and --outFilterMismatchNmax 3. PCR duplicates were identified by their UMI and removed using a custom script.

To obtain the spike-in normalization factor, rRNA-filtered reads > 20 nt were mapped to mouse mitochondrial mRNAs using bowtie1 and allowing one 5' and zero internal mismatches. After removing PCR duplicates the number of reads mapped was used as the spike-in normalization factor. This strategy results in roughly the fraction of spike-in reads expected from volume:volume fraction of mouse lysate added (5 or 10%), and ≤ 0.1% in samples without added mouse lysate (see **Table S1**).

The offset of the A site from the 3' end of reads was calibrated using start codons of *MT-ATP6*, *MT-CO3*, and *MT-ND4*, the only CDSs with significant 5' UTRs. Additionally, periodicity across CDSs was calculated for read 3' ends for each individual length. By determining the most common subcodon position for 3' ends of each read size across CDSs, the precise A site offset could be set for each experiment. Offsets from the 3' end of 30:[-13], 31:[-14], 32:[-15], 33:[-15], 34:[-15 or -16] were typically used. Soft-clipped nucleotides were omitted unless they were 3' As and thus appear to be part of a poly(A) tail. Shorter read lengths have ambiguous offsets because there is not consistency between whether 5' or 3' ends stay constant at a given mitoribosome position, and were excluded. A-site transformation was applied to mitochondrial mRNA-mapping reads using a custom script.

Mitoribosome profiling datasets for BJ fibroblasts ('Fibro-Rooijers') (*50*), additional HEK293T data ('HEK_Pearce') (*51-52*), and HCT116 cells (HCT116_Li-1, -2) (*53*) were downloaded from GEO (GSE48933 and GSE133315), ArrayExpress (E-MTAB-5519), and the SRA (SRR10491343 and SRR10491342).

Relative ribosome occupancies for codons were computed by taking the ratio of the A-site transformed ribosome density in a 3-nt window at the codon to the overall density in the coding sequence.

<u>Cytoribosome profiling data analysis</u>

Read trimming, filtering, alignment, and PCR duplicate removal was performed identically to mitoribosome profiling data analysis except that filtered reads were aligned to the human genome reference alone without the mouse reference.

Cytoribosome profiling datasets for HeLa S3 cells and fibroblasts from (*54*) and (*55*) were downloaded from GEO (GSE69906 and GSE115162).

<u>RNA-seq data analysis</u>

Read trimming, filtering, alignment, and PCR duplicate removal was performed identically to mitoribosome profiling data analysis except that default STAR mismatch parameters were used.

<u>OXPHOS subunit expression analysis</u>

Relative synthesis for OXPHOS genes was calculated as length- and library size-normalized read counts. For mitochondria-encoded OXPHOS subunit relative synthesis, mitoribosome profiling read counts were summed across genes using Rsubread featureCounts (*56*) with parameters readShiftType = "upstream", readShiftSize = 14, read2pos = 3, allowMultiOverlap = TRUE, useMetaFeatures = TRUE, countMultiMappingReads = TRUE, strandSpecific = 1, GTF.featureType='CDS'. Importantly, the GTF annotation file was modified to exclude the first 6 codons of all genes (which are variable depending on whether there is a 5' UTR) and all overlapping regions. Multimapping reads are included because nuclear mitochondrial DNA (NUMT) exists in the nuclear genome but does not appear to be the source of reads in mitoribosome profiling libraries as NUMT loci do not have uniquely-mapping reads. For mitochondrial genes, relative synthesis is expressed as transcripts per ten thousand (tp10k) (RPK/sum(mitochondrial protein-coding gene RPK)*10000). Note relative values are identical regardless of the normalization value.

For nuclear DNA-encoded OXPHOS subunit relative synthesis, cytoribosome profiling read counts were summed across protein-coding genes using featureCounts with parameters readShiftType = "downstream", readShiftSize = 17, read2pos = 5, allowMultiOverlap = TRUE, useMetaFeatures = TRUE, countMultiMappingReads = TRUE, strandSpecific = 1, GTF.featureType='CDS'. Multi-mapping reads are included because several OXPHOS genes have pseudogenes that are not a significant source of uniquely-mapping reads. For nuclear DNA-encoded genes, relative synthesis is expressed as reads per kb per million mapped reads (RPKM) (RPK/sum(reads mapping to nuclear protein-coding genes)*1000000). Importantly, several nuclear DNA-encoded OXPHOS subunits have paralogs that encode alternative isoforms. These isoforms do not have significant stretches of nucleotide sequence identity, but they do encode similar proteins that may be interchangeable depending on cell/tissue type. Therefore, isoform

RPKM counts are summed for the analyses in Figures 2, 4, and S4 (e.g. COX4I1 and COX4I2 become COX4I1/2). Of note, relative synthesis values for OXPHOS complex subunits computed with this method are highly correlated (r=0.96, data not shown) with relative synthesis rate values computed from the same data using a masking approach (*57*), except that here we recover values for seven additional subunits.

For both mtDNA- and nDNA-encoded OXPHOS subunit relative RNA abundance, RNA-seq read counts were summed across protein-coding genes using featureCounts with parameters allowMultiOverlap = TRUE, minOverlap=22, countMultiMappingReads = TRUE, GTF.featureType='CDS'. The GTF annotation file was modified to exclude overlapping regions in mitochondrial genes. 'CDS' was chosen as the feature to count instead of 'exon' for two reasons. (1) UTR annotation is less reliable than CDS annotation and (2) unexpressed isoforms with long UTRs artificially reduce apparent abundance. Library size normalizations were performed separately for mtDNA- and nDNA--encoded subunits resulting in relative abundance values expressed as tp10k and RPKM, respectively, as described above for ribosome profiling data.

GO-term enrichment analysis of differentially expressed genes

To determine differential expression of the $LRPPRC^{KO}$ mutant compared to the *LRPPRC* rescue cells, we performed a differential expression analysis using RNA-seq reads counted as described in 'OXPHOS subunit expression analysis' above. For differential expression analysis the R package DESeq2 (*58*) was used. Default parameters were used and the counts were normalized using the estimateSizeFactors function. For differential expression results, the method contrast "sampletype" was used. To determine significantly changed genes, genes were filtered for adjusted p-value < 0.05. Two sets were generated for GO-term enrichment analysis, genes with log2 changes >= 1 and with log2 changes <= -1.

For GO-term enrichment analysis the R package clusterProfiler (*59*) was used. The unfiltered results table was used to generate the universe gene list used to test against for enrichment. The filtered lists for either significantly up- or down-regulated genes were used for testing. GO term analysis was performed using the enrichGO function with the following parameters: keyType = "ENSEMBL", OrgDb = org.Hs.eg.db, ont = "BP", pAdjustMethod = "BH", pvalueCutoff = 0.05, qvalueCutoff = 0.05, readable = TRUE). For Figure 4C**,** the resulting GO terms were filtered for redundancy, using the simplify function with the following settings: cutoff = 0.2, by = "p.adjust", select_fun = min, measure = "Wang", semData = NULL. For visualization, the dotplot function was used with showCategory = 10. For Figure S5I GO-terms without filtering were used.

*MT-CO3* initiation analysis

The fraction of processed and unprocessed *MT-ATP8/6-CO3* transcript was calculated from RNA-seq data using a custom python script. Reads with their 5' end precisely at *MT-CO3* start were counted as coming from processed transcripts, and reads that span the junction were counted as coming from unprocessed transcripts. The fraction of *MT-CO3* translation initiation on processed and unprocessed *MT-ATP8/6-CO3* transcript was calculated from mitoribosome profiling data using a custom R script. Reads with their 5' end precisely at *MT-CO3* start (vertical arrowhead in Figure 1G) were counted as coming from initiation on processed *MT-CO3*. Reads that span the junction could result from either *MT-CO3* initiation or *MT-ATP6* termination. Therefore, to count only instances of initiation on unprocessed *MT-CO3*, reads with their 5' ends highlighted by the diagonal arrowhead in Figure 1G were summed.

Ad hoc scoring

Start codons were assigned scores based on mitoribosome profiling data as described in Figure S3C using a custom R script. Briefly, the genome sequence was split into three frame references and A-site reads falling on the second (middle) position of each were assigned to that reference. Each start and stop codon was also assigned to a reference, as were known genes. Scores were assigned to each AUG, AUU, AUA, and GUG codon in three parts where the final score is the sum of: 1) Summed fold-change in P-site in-frame reads (inferred from A-site reads one codon downstream) compared to one codon upstream and one codon downstream. 2) Average percent of in-frame reads 20 codons downstream minus average percent of in-frame reads 16 codons upstream. 3) Total number of reads (in all frame references) five codons downstream divided by total number of reads five codons upstream. This final value was log-transformed then doubled to keep its contribution commensurate with the other values. For score frequency histograms shown in Figures 1J and S3C, start codons within the reading frame of known ORFs were excluded and rRNA- and tRNA-filtered heavy strand data was used, resulting in no scores for start codons in rRNA or tRNA genes, nor on the light strand (e.g. *MT-ND6*).

SNV analysis

Alternative start codons resulting from homoplasmic variants (*36*) at ATG codons were counted using a custom R script. Because *MT-ND5-dORF* is encoded by the same DNA (but on the opposite strand) as *MT-ND6*, we limited our analysis to other ATGs in the identical orientation antisense to known genes (see **Figure 1K,L**). ATT, ATA, and GTG are counted as alternative start codons, but not ACG, which is capable of translation initiation in vitro (*37*), but has not been found to be used in vivo in human mitochondria.

Fig. S1.

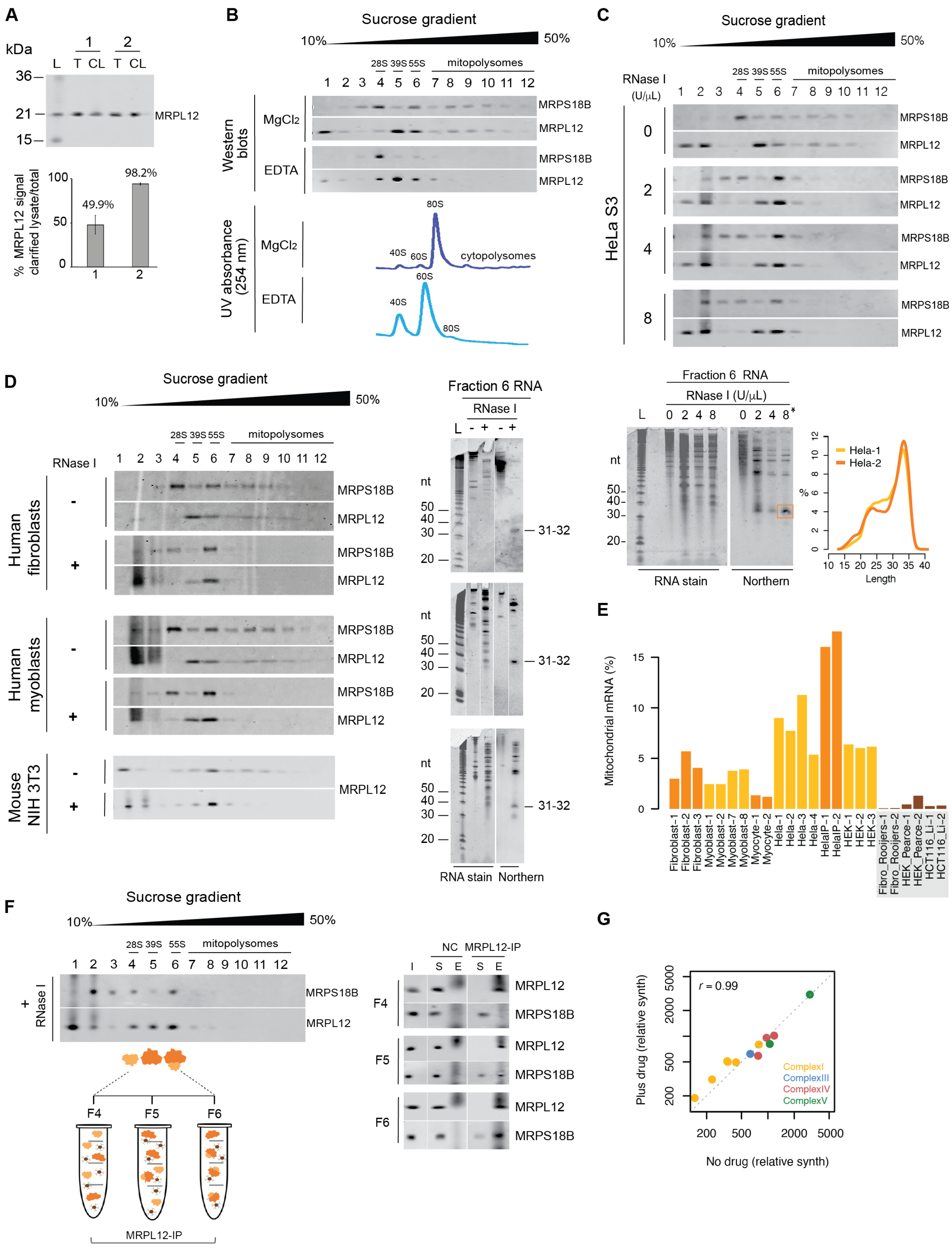
A
kDa
L T CL T CL
MRPL12
% MRPL12 signal clarified lysate/total
49.9%
98.2%
B
Sucrose gradient
10%
50%
28S 39S 55S
mitopolysomes
Western blots
MgCl2
EDTA
MRPS18B
MRPL12
UV absorbance (254 nm)
40S 60S 80S
cytopolysomes
C
HeLa S3
RNase I (U/μL)
D
RNase I
Human fibroblasts
Human myoblasts
Mouse NIH 3T3
Fraction 6 RNA
nt
31-32
RNA stain
Northern
Fraction 6 RNA
RNase I (U/μL)
Hela-1
Hela-2
Length
E
Mitochondrial mRNA (%)
Fibroblast-1
Fibroblast-2
Fibroblast-3
Myoblast-1
Myoblast-2
Myoblast-7
Myoblast-8
Myocyte-1
Myocyte-2
Hela-1
Hela-2
Hela-3
Hela-4
HelaIP-1
HelaIP-2
HEK-1
HEK-2
HEK-3
Fibro_Rooijers-1
Fibro_Rooijers-2
HEK_Pearce-1
HEK_Pearce-2
HCT116_Li-1
HCT116_Li-2
F
Sucrose gradient
RNase I
NC
MRPL12-IP
I S E S E
F4
F5
F6
MRPL12-IP
G
r = 0.99
ComplexI
ComplexIII
ComplexIV
ComplexV
Plus drug (relative synth)
No drug (relative synth)

**Fig. S1. Optimization and adaptability of mitoribosome profiling.** (**A**) Mitoribosome extraction efficiency after employing previously established (*50*) solubilization conditions (1), and our optimized lysis buffer (2). HeLa S3 cells were lysed with either lysis composition, followed by clarification of the lysates. Equal volumes of total (T) and clarified lysates (CL) were subjected to western blots against a mitoribosome subunit antibody (MRPL12). (L) indicates the protein marker used in kilodaltons (kDa). Quantification from two experiments of the total and post-clarification signals revealed the optimized conditions allow for the recovery of >98% of mitoribosomes, compared to only 50% for the existing protocols. Error bars represent the range between two replicates. (**B**) Sedimentation of mito- and cytoribosomes in 10-50% linear sucrose gradients. HeLa S3 cells were lysed in the presence of either $MgCl_2$ or EDTA. Clarified lysates were loaded onto linear 10-50% sucrose gradients. Detection of mitoribosome (top panel) in each fraction was achieved by western blots against proteins of the large (MRPL12) and small subunit (MRPS18B). We followed the sedimentation of cytoribosomes by their UV absorbance at 254 nM (bottom panel). (**C**) Titration of RNase I to generate mitoribosome footprints. Increasing concentrations (0, 2, 4 and 8 Units/µL) of the nuclease were used to digest 450 µl of HeLa S3 lysate at room temperature for 30 minutes. Digested lysates were further clarified and loaded onto 10-50% linear sucrose gradients. Detection of mitoribosomes in each fraction (top panel) was achieved by western blots against proteins of the large (MRPL12) and small subunit (MRPS18B). Protected mitoribosome footprints from the mitomonosome (Fraction 6 RNA) were further analyzed by northern blot (bottom panel), showing a robust and compact footprint for the highest RNase I amount used (8 Units/µL), denoted with an (*) as the concentration used in subsequent sample generation. The fragment length distribution from this sample (marked inside the box), and a replicate of reads mapping to mitochondrial mRNAs are shown to the right. (**D**) Mitoribosome isolation conditions were readily adaptable to other cell lines. As shown in **(C)** for HeLa S3 cells, digested lysates from human fibroblasts, human myoblasts and mouse NIH3T3 were used to generate and isolate mitoribosome footprints following the sedimentation of the mitomonosome (fraction 6 RNA) by western blots of large (MRPL12) and small (MRPS18B) mitoribosome subunits (left panels). Footprint size was assessed by northern blot (right panels). (**E**) Percentage of reads in each mitoribosome profiling sample that map to mitochondrial mRNAs. Gray box highlights published WT, untreated samples from BJ fibroblasts (*50*) (Fibro_Rooijers-1, -2), HEK293T cells (*52*) (HEK_Pearce-1) and (*51*) (HEK_Pearce-2), and HCT116 cells (*53*) (HCT116_Li-1, -2) analyzed in parallel. Full sample compositions are shown in Table S1. (**F**) Mitoribosome immunoprecipitation. RNase I-digested HeLa S3 lysates were clarified and loaded onto 10-50 % sucrose gradients (left panel). Fractions 4, 5 and 6 were incubated with either MRPL12-conjugated DynaBeads or naked DynaBeads (NC) as negative control. We analyzed subsequent input (I), supernatant (S), and elution products (E) using western blots (right panel), observing the highest amount of the small subunit immunoprecipitated in fraction 6. (**G**) Comparison of relative synthesis rate values (tp10k) for mtDNA-encoded OXPHOS subunits in cells exposed or not to ribosome inhibitors. Pearson correlation, r. Before lysis, cells were treated with cycloheximide (100 μg/mL) and chloramphenicol (100 μg/mL) for 15 and 5 minutes, respectively, to stall cytosolic and mitochondrial translation. After treatment, cells were washed with 1X PBS, pH 7.4 and subjected to lysis using our mitoribosome profiling conditions.

**Fig. S2.**

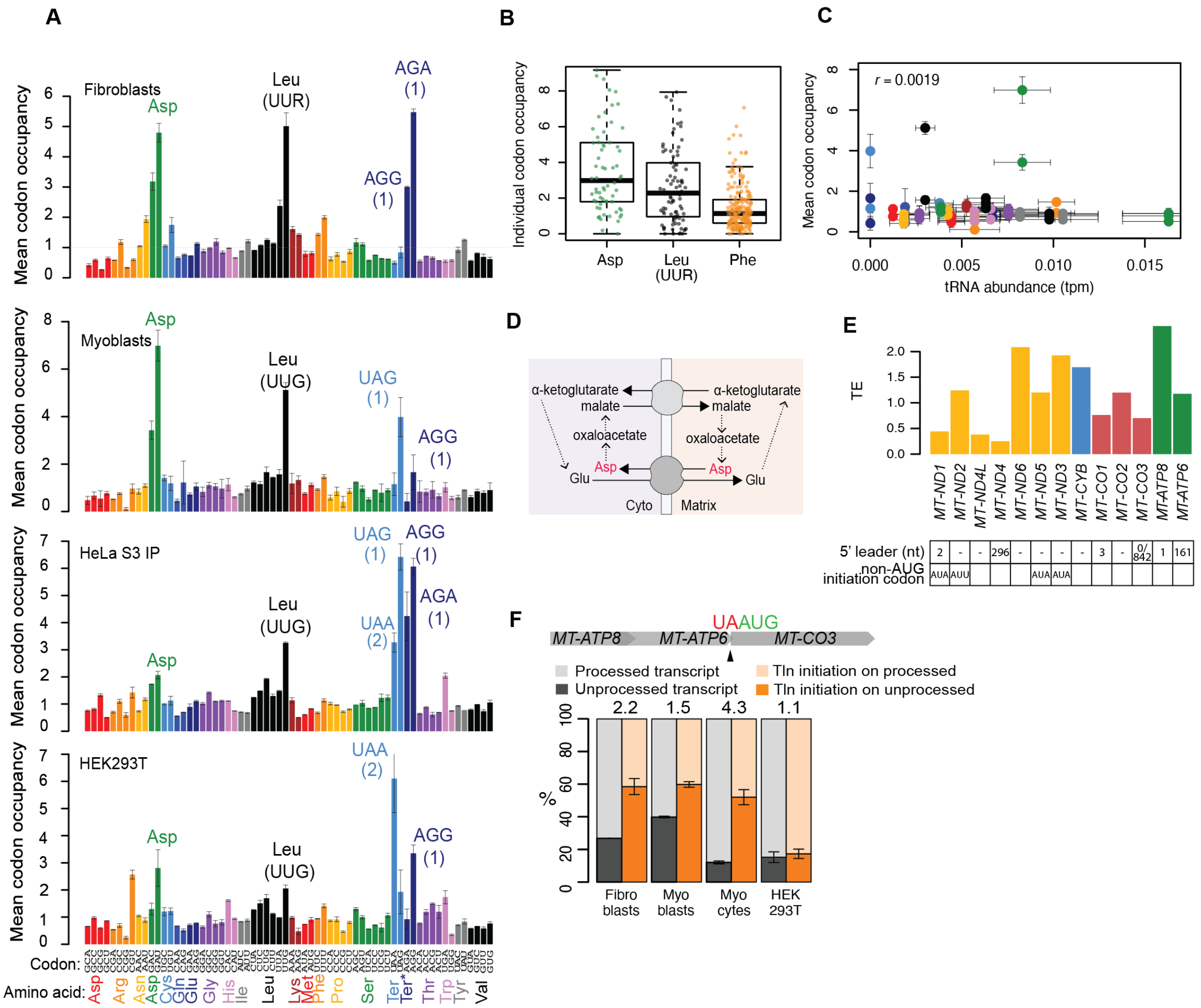
A
B
C
D
E
F
Fibroblasts
Myoblasts
HeLa S3 IP
HEK293T
Mean codon occupancy
Codon:
Amino acid:
Individual codon occupancy
tRNA abundance (tpm)
r = 0.0019
α-ketoglutarate
malate
oxaloacetate
Asp
Glu
Cyto
Matrix
TE
5' leader (nt)
non-AUG initiation codon
MT-ATP8
MT-ATP6
MT-CO3
UAAUG
Processed transcript
Unprocessed transcript
TIn initiation on processed
TIn initiation on unprocessed
Fibro blasts
Myo blasts
Myo cytes
HEK 293T

**Figure S2**. **Insights into mitochondrial translation.** (**A**) Mitoribosome occupancy across codons. Asp, Leu (UUR), and termination codons are labeled. AGA and AGG Ter* putative termination codons are each present only once (indicated in parentheses), at the ends of *MT-CO1* and *MT-ND6*, respectively. R: purine (A or G). Codons with occupancy of >3x expected are labeled. Error bars show range across replicates. (**B)** Distribution of occupancies across individual Asp, Leu (UUR) and Phe codons in fibroblasts. Dotted line highlights expected occupancy in the absence of pausing. (**C**) Mean codon occupancy in myoblasts compared to tRNA abundance in myoblasts (*28*). Color coding is the same as in (A). Error bars for mean codon occupancy show range across 4 replicates. Error bars for tRNA abundance show standard deviation across 3 replicates. Pearson correlation, r, is excluding termination codons (shown at tRNA abundance = 0). (**D**) Malate aspartate shuttle scheme. (**E**) Translation efficiency (TE = relative synthesis values/relative RNA abundance values) of each mitochondrial OXPHOS transcript in fibroblasts. Transcript 5' end characteristics are listed below plot. (**F**) Percentage of processed and unprocessed *MT-ATP8/6-CO3* transcript at site indicated by arrowhead and of translation (Tln) initiation on processed and unprocessed transcript. Numbers at top of bars indicate the fold-preference for translation initiation on unprocessed vs processed transcripts. Error bars show range across replicates.

### Figure S3

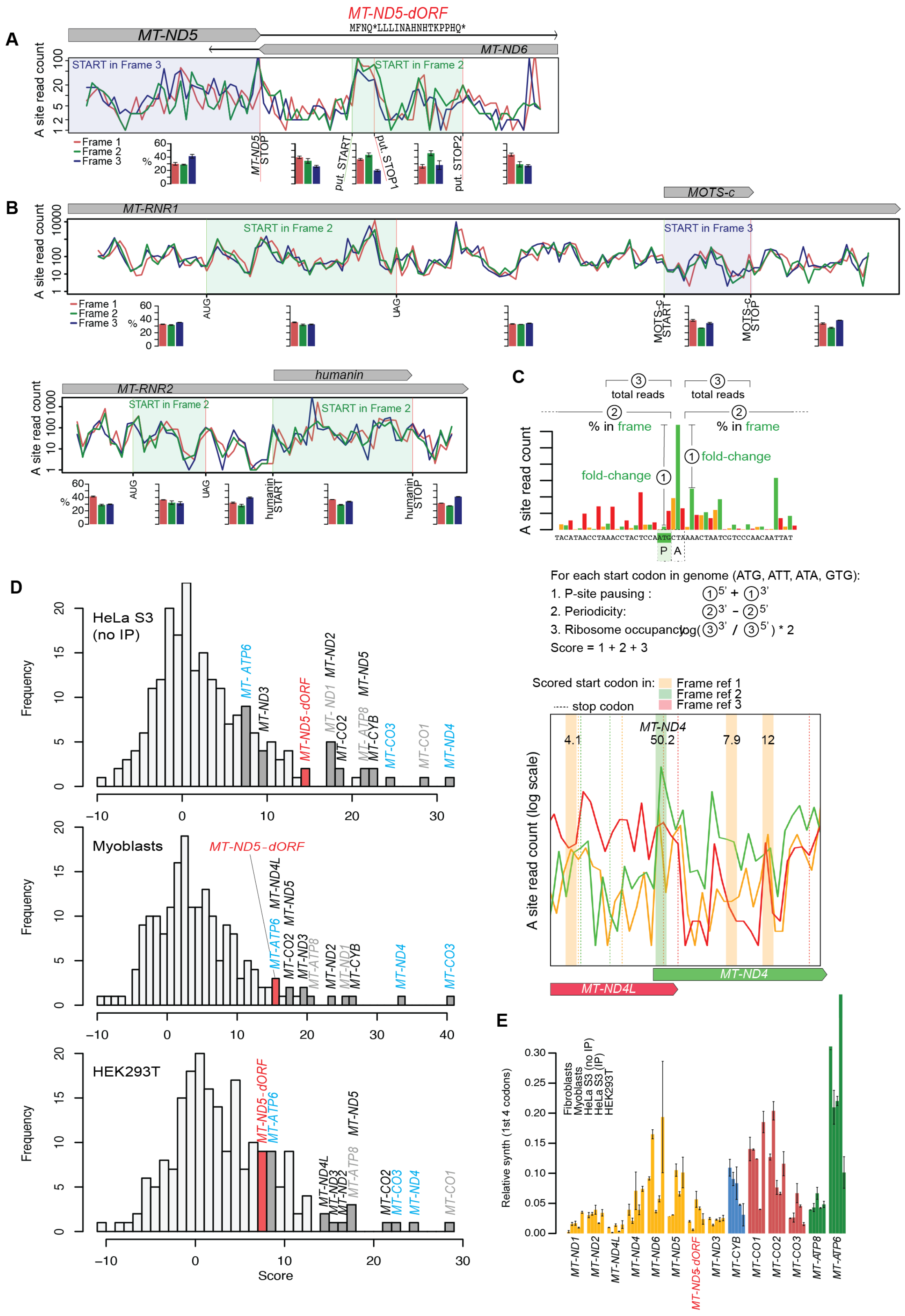

A
MT-ND5-dORF
MFNQ*LLLINAHNHTKPPHQ*
MT-ND5
MT-ND6
START in Frame 3
START in Frame 2
A site read count
Frame 1
Frame 2
Frame 3
MT-ND5 STOP
put. START
put. STOP1
put. STOP2
B
MOTS-c
MT-RNR1
START in Frame 2
START in Frame 3
AUG
UAG
MOTS-c START
MOTS-c STOP
humanin
MT-RNR2
humanin START
humanin STOP
C
total reads
% in frame
fold-change
A site read count
For each start codon in genome (ATG, ATT, ATA, GTG):
1. P-site pausing :
2. Periodicity:
3. Ribosome occupancy
Score = 1 + 2 + 3
Scored start codon in:
Frame ref 1
Frame ref 2
Frame ref 3
stop codon
MT-ND4
A site read count (log scale)
MT-ND4L
D
HeLa S3 (no IP)
Myoblasts
HEK293T
Frequency
Score
E
Fibroblasts
Myoblasts
HeLa S3 (no IP)
HeLa S3 (IP)
HEK293T
Relative synth (1st 4 codons)

**Figure S3**. **Evidence for the translation of a novel mitochondrial open reading frame**. (**A**) Stacked frame plot highlighting number of A-site transformed reads in each subcodon position as in Figure 1I, but for HeLa S3 cells (IP data). Mitoribosome profiling data are from two replicates, summed. Bar plots show the average percentage of reads in each subcodon position for regions indicated, with error bars showing range between the replicates summed above. (**B**) Stacked frames plots across mitochondrial rRNA genes *MT-RNR1* and *MT-RNR2* in the regions of *MOTS-c* and *humanin*, respectively. One other short, unannotated open reading frame is shown in each region. Data is from two fibroblast replicates, summed. Bar plots show the average percentage of reads in each subcodon position for regions indicated, with error bars showing range between the replicates summed above. (**C**) Ad hoc scoring strategy to identify likelihood of translation initiation at each start codon. Top panel: Details of scoring computation. Windows used in {2} are 16 codons upstream and 20 codons downstream. Window used in {3} is five codons. See methods for more details. Bottom panel: example output including region of raw data shown above. Wide vertical lines show putative start codons with their scores, with the color indicating their frame. Dotted vertical lines show putative stop codons, with the color indicating their frame. The number at top are the ad hoc scores. (**D**) Ad hoc score distributions, as in Figure IJ, for additional cell lines. Gene names in black: known genes with no leader; gray: known genes with 1 to 3 nt leader; cyan: known genes with long 5' UTR. (**E**) Estimated relative synthesis of *MT-ND5-dORF* compared to other mtDNAencoded genes. A-site transformed reads were summed across the first four codons with signal for each gene and normalized by the total number of such reads. For genes with 5' UTRs (*MT-ATP6*, *MT-ND4*, *MT-CO3*, *MT-ND5-dORF*) this includes codons 2-5, where codon 5 is a stop codon for *MT-ND5-dORF*. For genes with a very short or no leader (all others) this includes codons 6-9. For *MT-ATP6*, the signal is confounded by reads from *MT-ATP8*, which overlaps the first 15 codons. Samples are plotted in the order listed and error bars show range across replicates.

**Fig. S4.**

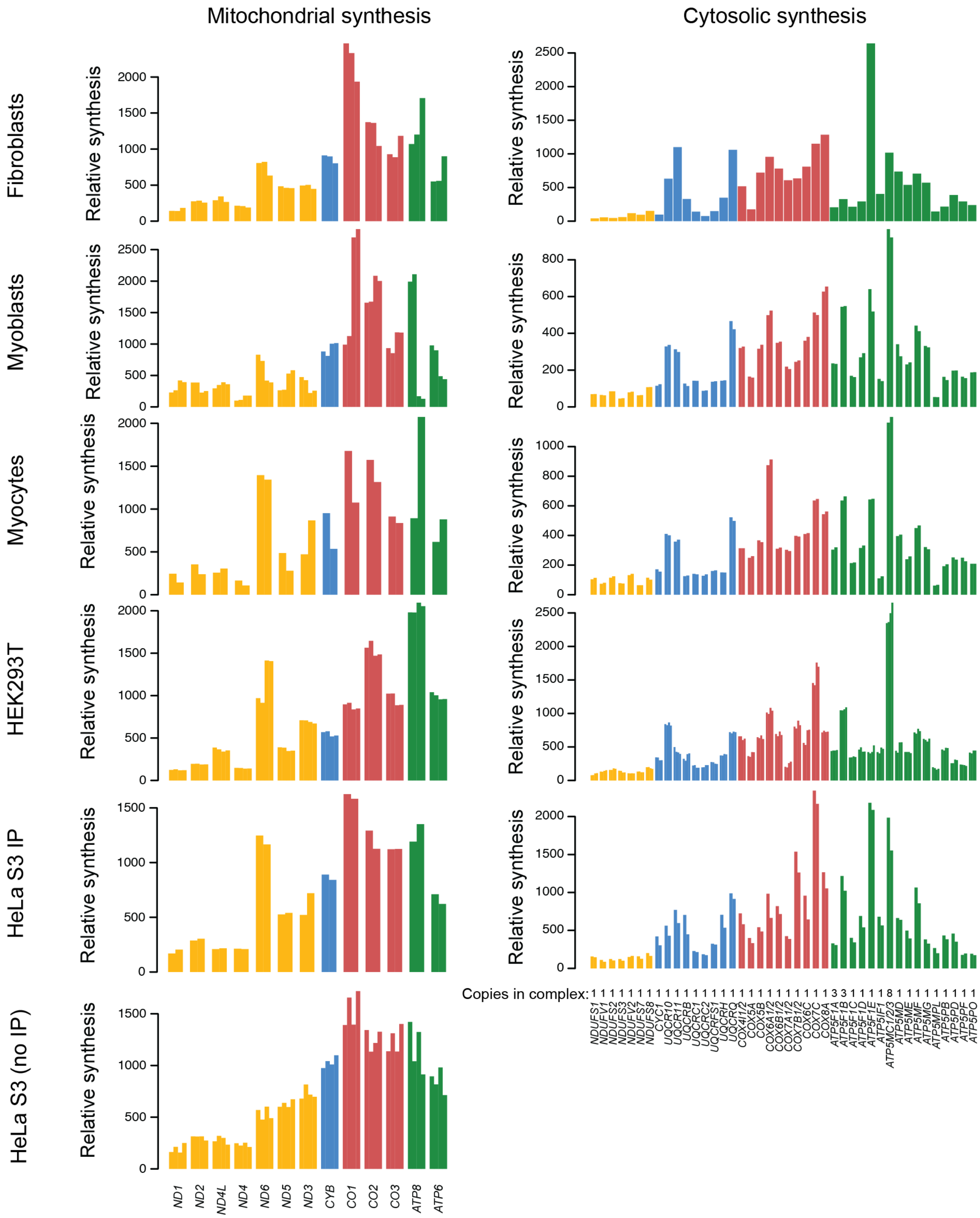
Mitochondrial synthesis
Cytosolic synthesis
Fibroblasts
Myoblasts
Myocytes
HEK293T
HeLa S3 IP
HeLa S3 (no IP)
Relative synthesis
Copies in complex: 1 1 1 1 1 1 1 1 1 1 1 1 1 1 1 1 1 1 1 1 1 1 1 1 1 1 3 3 1 1 1 1 8 1 1 1 1 1 1 1 1 1
NDUFS1 NDUFV1 NDUFS2 NDUFS3 NDUFV2 NDUFS7 NDUFS8 CYC1 UQCR10 UQCR11 UQCRB UQCRC1 UQCRC2 UQCRFS1 UQCRH UQCRQ COX4I1/2 COX5A COX5B COX6A1/2 COX6B1/2 COX7A1/2 COX7B1/2 COX6C COX7C COX8A ATP5F1A ATP5F1B ATP5F1C ATP5F1D ATP5F1E ATP5IF1 ATP5MC1/2/3 ATP5MD ATP5ME ATP5MF ATP5MG ATP5MPL ATP5PB ATP5PD ATP5PF ATP5PO
ND1 ND2 ND4L ND4 ND6 ND5 ND3 CYB CO1 CO2 CO3 ATP8 ATP6

**Figure S4**. **Correlated synthesis of mtDNA- and nDNA-encoded OXPHOS subunits.** All data included in Figure 2B, C. Each replicate is shown as an individual bar, values given in Table S2. Cytosolic synthesis values for fibroblasts and HeLa S3 cells were calculated from cytosolic ribosome profiling data from (*55*) and (*54*), respectively. Relative synthesis is measured in tp10k for mitochondrial synthesis and RPKM for cytosolic synthesis (see Methods for details and note that mitochondrial and cytosolic synthesis absolute values cannot be compared across compartments).

### Fig. S5.

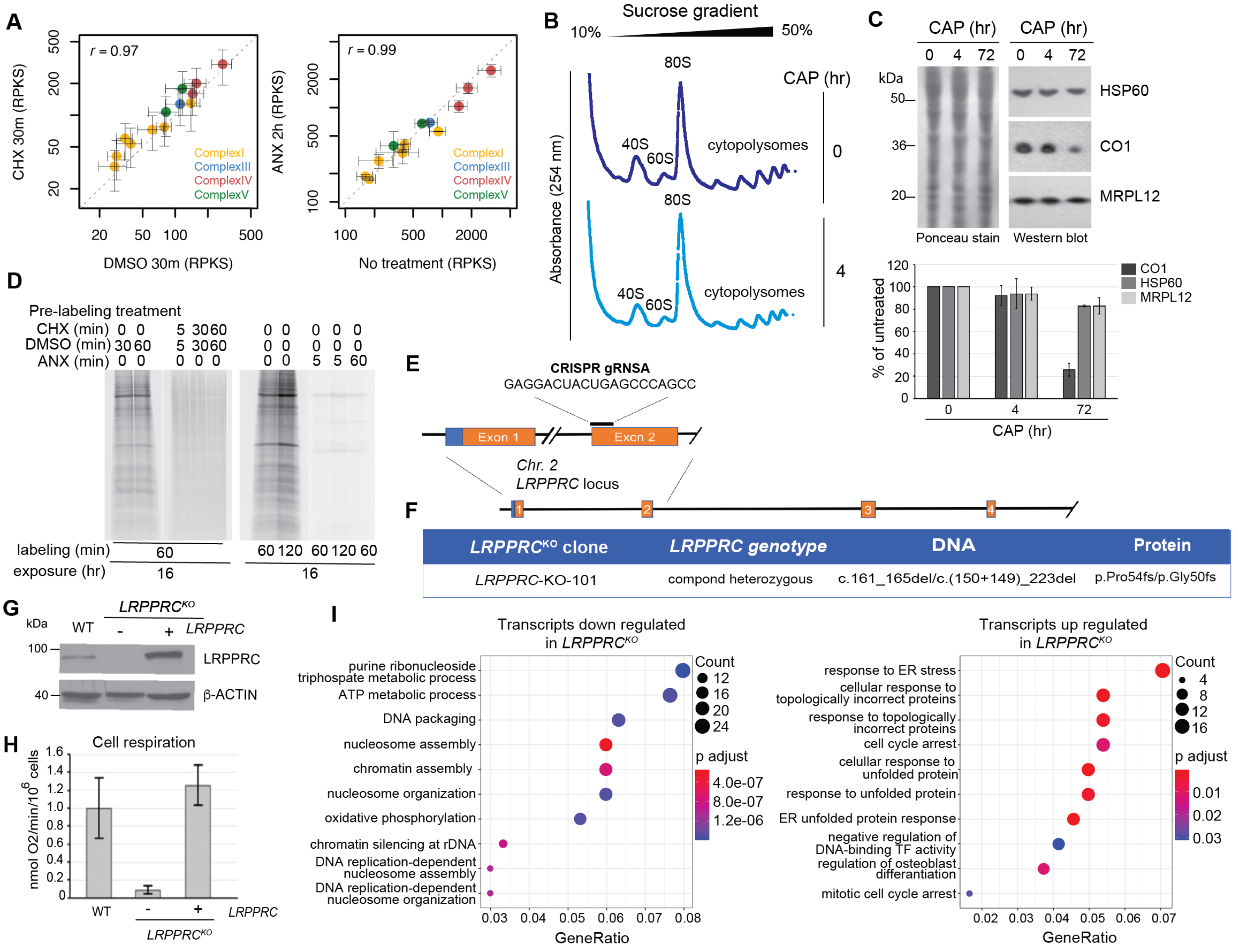

A
r = 0.97
CHX 30m (RPKS)
DMSO 30m (RPKS)
ComplexI
ComplexIII
ComplexIV
ComplexV
r = 0.99
ANX 2h (RPKS)
No treatment (RPKS)
B
Sucrose gradient
10%
50%
Absorbance (254 nm)
CAP (hr)
80S
40S
60S
cytopolysomes
0
4
C
CAP (hr)
kDa
HSP60
CO1
MRPL12
Ponceau stain
Western blot
% of untreated
CAP (hr)
D
Pre-labeling treatment
CHX (min)
DMSO (min)
ANX (min)
labeling (min)
exposure (hr)
E
CRISPR gRNSA
GAGGACUACUGAGCCCAGCC
Exon 1
Exon 2
Chr. 2
LRPPRC locus
F
LRPPRC^KO clone | LRPPRC genotype | DNA | Protein
LRPPRC-KO-101 | compond heterozygous | c.161_165del/c.(150+149)_223del | p.Pro54fs/p.Gly50fs
G
LRPPRC^KO
WT
LRPPRC
β-ACTIN
H
Cell respiration
nmol O2/min/10^6 cells
I
Transcripts down regulated in LRPPRC^KO
purine ribonucleoside triphospate metabolic process
ATP metabolic process
DNA packaging
nucleosome assembly
chromatin assembly
nucleosome organization
oxidative phosphorylation
chromatin silencing at rDNA
DNA replication-dependent nucleosome assembly
DNA replication-dependent nucleosome organization
Count
p adjust
GeneRatio
Transcripts up regulated in LRPPRC^KO
response to ER stress
cellular response to topologically incorrect proteins
response to topologically incorrect proteins
cell cycle arrest
cellular response to unfolded protein
response to unfolded protein
ER unfolded protein response
negative regulation of DNA-binding TF activity
regulation of osteoblast differantiation
mitotic cell cycle arrest

**Figure S5. Transcription perturbations and ablation of *LRPPRC*.** (**A**) Comparison of RPKS (mouse spike-in normalized reads per kb) values for mtDNA-encoded OXPHOS subunits with and without 30 minutes of cycloheximide (CHX) (100 µg/mL) or 2 hours of anisomycin (ANX) treatment (100 µg/mL) to inhibit cytosolic translation. (**B**) Sedimentation of cytosolic polysomes in 10-50% linear sucrose gradients. HEK293T cells were treated with mitochondrial ribosome inhibitor chloramphenicol (CAP) for the indicated times in hours (hr). Cells were subsequently lysed and loaded onto gradients as explained in the methods section for cytoribosome profiling. Cytoribosomes were detected by their UV absorbance at 254 nm. (**C**) Western blot analysis of HEK293T cells treated with chloramphenicol (CAP) for the indicated times (top panel). Protein marker molecular weights are indicated in kDa. Quantification shows the signal percentage of untreated cells normalized by Ponceau stain (bottom panel). (**D**) *In vivo* labeling of translation products. Human fibroblasts newly synthesized products were metabolically labeled with 35S methionine/cysteine, after treatment with either cycloheximide (CHX), anisomycin (ANX) or DMSO control, for the indicated times. Signals were exposed to a screen for 16 hrs, as explained in Methods. (**E**) Schematic showing the location of the target sites in the *LRPPRC* locus for the CRISPR guide RNA used to knock out the gene. (**F**) Table detailing the genotype of the generated HEK293T cell line carrying edited *LRPPRC* alleles. (**G**) Immunoblot analysis of LRPPRC steady-state levels in WT and *LRPPRC*$^{KO}$ cells and *LRPPRC*$^{KO}$ cells reconstituted with recombinant *LRPPRC*. An antibody against β-actin was used as the loading control. Protein marker molecular weights are indicated in kDa. (**H**) Endogenous cell respiration in WT, *LRPPRC*$^{KO}$, and WT rescue cell lines measured polarographically. (**I**) GO-term enrichment analysis of significantly (adjusted p-value < 0.05) differentially expressed genes that are at least two-fold decreased or increased in the *LRPPRC*$^{KO}$ cells compared to the WT rescue cell line. p adjust = adjusted p-value, TF = transcription factor.

**Table S1.**
Provided as TableS1.xlsx

**Table S1. Library composition and quality characteristics for wild type, untreated samples.** Input counts are the number of reads with adapter found that are ≥ 11 nt long after adapter and UMI trimming. The size range used for A site transformation was chosen based on the peak of RPF read length distribution (e.g. full length reads) and unambiguous offset determination. Periodicity is the percent of reads in the size range listed that have A sites on the second subcodon position. For coverage, the first six codons of each known ORF are excluded because full-length reads do not align to these positions for the majority of genes since they do not have 5' UTRs. A codon is considered covered if it has at least one read on any of the three subcodon positions. Notably, a lack of coverage can make periodicity appear more substantial than it is. To account for this we use coverage-adjusted periodicity which is the unadjusted periodicity multiplied by the fraction of codons covered. Published datasets not generated in this study, but analyzed in parallel, are highlighted in grey. Important metrics of library quality are highlighted in orange.

**Table S2.**
Provided as TableS2.xlsx

**Table S2. Relative synthesis values for OXPHOS subunits.** Mitoribosome profiling, cytoribosome profiling, and RNA-seq relative synthesis values used in Figures 2, 4, and S4. mtDNA-encoded subunit synthesis is expressed in transcripts per 10,000, and nDNA-encoded subunit synthesis is expressed in RPKM as described in Methods.

**Table S3.**
Provided as TableS3.xlsx

**Table S3. Differential expressed (DE) genes in *LRPPRC*$^{KO}$.** Log2 fold expression changes from *LRPPRC*$^{KO}$ cells compared to the *LRPPRC* rescue cells used for the GO enrichment analysis.

baseMean = mean of normalized counts for all samples, Log2FoldChange = log2 fold change (MLE): sampletype KO vs KOplWT, lcfSE =standard error: sampletype KO vs KOplWT, stat = Wald statistic: sampletype KO vs KOplWT, p-value = Wald test p-value: sampletype KO vs KOplWT, p adjust = BH (Benjamini, Hochberg) adjusted p-values. Shown are the counts used for each strain and replicate (n=2). KO = *LRPPRC*$^{KO}$, KOplWT = *LRPPRC* rescue. Significant column marks significantly changed genes (NA = not available, Not Sig = not significant, FDR < 0.05 = significant).

**Table S4.**
Provided as  as TableS4.xlsx

**Table S4: GO terms enriched for differential expressed genes in *LRPPRC*$^{KO}$.** Full list of GO terms enriched for at least 2 fold up or down regulated genes used in Figures 4C and S5I. ID = GO term ID, Description = GO term description, GeneRatio = number of genes with GO term devided by total number of tested genes, BgRatio = total number of genes with GO term devided by universe list (all genes used for DESeq2 analysis), pvalue = calculated p value using hypergeometric distribution, p adjust = adjusted p value using BH method, qvalue = false discovery rate, GeneID = list of genes that share the respective GO term, Count = number of genes that share the respective GO term.
